# Age-dependent topoisomerase I depletion alters recruitment of rDNA silencing complexes

**DOI:** 10.1101/2025.07.29.667507

**Authors:** Lindsey N. Power, Natalia Zawrotna, Manikarna Dinda, Abigail E. Weir, Bishal P. Paudel, Oshil Ghimire, Karolina Kisiel, Christopher T. Letai, Kevin A. Janes, Jeffrey S. Smith

**Affiliations:** Department of Biochemistry and Molecular Genetics, University of Virginia School of Medicine, Charlottesville VA; Department of Medical Chemistry, Medical University of Gdansk, Gdansk Poland; Department of Biomedical Engineering, University of Virginia School of Medicine, Charlottesville VA; Department of Microbiology, Immunology, and Cancer, University of Virginia School of Medicine, Charlottesville VA; College of Arts and Sciences, University of Virginia, Charlottesville VA; College of Health Science, Physician Assistant Program Midwestern University, Downers Grove IL

## Abstract

Genomic instability and loss of proteostasis are two of the primary Hallmarks of Aging. Although these hallmarks are well-defined in the literature, the mechanisms that drive genomic instability and loss of proteostasis as cells age are still largely unknown. Using budding yeast replicative lifespan as a model for aging in actively dividing cells, we identified nuclear proteins that were depleted in the earliest stages of aging. We found that many age-depleted proteins were involved in ribosome biogenesis, specifically in ribosome processing, or in maintenance of chromatin stability. We focused on topoisomerase I (Top1) as a novel age-depleted nuclear protein and found that its depletion in the early stages of aging was not a result of transcriptional changes or changes in protein turnover. Despite the stark depletion of Top1 in early aging, rescue of this age-dependent depletion was actually harmful to replicative lifespan. We found that Top1, when overexpressed, disrupts the stoichiometry of the RENT complex by pulling Sir2 away from the ribosomal DNA (rDNA), a phenotype which is further enhanced when the overexpressed Top1 is catalytically dead. Loss of Sir2 from the rDNA via the overexpression of catalytically dead Top1 decreases RNA Pol II silencing of a reporter gene inside or adjacent to the rDNA, consistent with the lifespan defect. Finally, we found that the catalytic activity of Top1 plays an important role in the establishment of rDNA silencing, raising the possibility that rDNA secondary structure/DNA topology is important for RNA Pol I-dependent spreading of silent chromatin across the rDNA locus.

## INTRODUCTION

Aging across eukaryotes involves a series of declining cellular functions known commonly as the “Hallmarks of Aging”. The molecular pathways involved in some of these hallmarks, such as genome instability and loss of proteostasis, are highly conserved from yeast to humans (1, 2). Due to this high level of conservation, replicative aging in *Saccharomyces cerevisiae* (budding yeast) has long been an established model for studying the molecular mechanisms of aging in dividing cells since the discovery that yeast have a finite lifespan (2). The replicative lifespan (RLS) of budding yeast is quantified as the number of times a mother cell buds before death (2). Importantly, the daughter cells bud asymmetrically from the aging mother cell and have full replicative potential, so RLS is thought to be a model for aging in dividing cells of higher eukaryotes, such as stem cells (3).

A notable example of an aging-relevant discovery from yeast was the identification of Sir2 and other sirtuin family members as NAD^+^-dependent histone/protein deacetylases (HDACs) (4–6). Sir2 is the catalytic subunit of two HDAC complexes that function in heterochromatic gene silencing (7, 8). The first Sir2 HDAC complex, known as the SIR complex, consists of Sir2, Sir3, and Sir4, and is responsible for the establishment and maintenance of silencing at telomeres and the silent mating type loci (*HML* and *HMR*) (9–11). The second Sir2 HDAC complex is known as RENT (<u>RE</u>gulator of <u>N</u>ucleolar silencing and <u>T</u>elophase exit) and consists of Sir2, Net1, and Cdc14 subunits (12, 13). RENT is recruited to the IGS1 and IGS2 intergenic spacers of the rDNA locus where it silences RNA Pol II-mediated transcription of several lncRNAs (11, 14–16), a function required to maintain stability of the tandem array (17, 18). Importantly, recruitment of the RENT complex to the rDNA to induce silencing is partially dependent on active RNA Pol I transcription of the 35S rRNA gene (19).

The rDNA array is a key modulator of replicative lifespan. Moderate overexpression of Sir2 enhances rDNA silencing and extends RLS (20, 21). Furthermore, the rDNA tandem array stability and array size have direct effects on RLS. For example, yeast mother cells accumulate extrachromosomal rDNA circles (ERCs) as they age due to unequal crossing over during DSB repair at the Fob1-mediated replication fork block (RFB) (22, 23). Destabilizing the array by deleting *SIR2* increases the number of ERCs and causes premature aging (20), but this increase in ERCs/reduced lifespan is partially rescued by deletion of *FOB1* (24), which suppresses ERC production. Additionally, yeast RLS correlates directly with rDNA copy number, as strains with greater rDNA copy number in the tandem array tend to have better longevity (25). Because rDNA stability is such an important modulator of lifespan, maintenance of rDNA regulatory and structural complexes such as RENT and others becomes critical as cells age.

Given that overall translational efficiency decreases globally in aged yeast cells (26), many important proteins may decline in abundance during replicative aging (26). A decline in certain protein levels can alter stoichiometry of multi-subunit protein complexes and have negative consequences for RLS. For example, altered stoichiometry of nuclear pore complexes has been linked to replicative aging (27). Additionally, the dysregulation of protein level homeostasis in aging yeast has been linked to genome instability, as cells with increased rDNA instability undergo aggregation of important rRNA binding proteins (28). Key rDNA silencing proteins like Sir2 are limiting within the cell, giving rise to competition between the rDNA locus and other regions of the genome where these proteins are functioning, such as telomeres (21). Potentially exacerbating this competition, Sir2 protein levels were previously shown to be significantly reduced in replicatively aging yeast cells (29).

It has also been demonstrated that the cohesin subunits Mcd1 and Scc1 are depleted during replicative aging and that forced depletion of Mcd1 shortens RLS (30, 31). Mcd1 is also depleted from the rDNA locus at the early stages of aging and redistributed to centromeres (30), most likely to retain the integrity of mitotic chromosome segregation. During the later stages of aging, as cells develop a general chromosome instability phenotype, any remaining Mcd1 is also depleted from centromeres, correlating with a general chromosome instability phenotype (30). Overexpression of *MCD1* from a doxycycline inducible promoter is sufficient to stabilize the rDNA and extend RLS (30), similar to the effect of increasing Sir2 expression (20). These studies led us to ask what other nuclear proteins are depleted during the early stages of replicative aging and thus important in maintaining rDNA stability.

Using a mini chemostat aging device (MAD) system (32), we performed a proteomic screen for nuclear proteins that significantly change in abundance during the early stages of replicative aging. We identified several important chromatin stability factors such as topoisomerases, helicases, and remodelers, as well as ribosome biogenesis factors that were significantly reduced. Focusing on topoisomerase I (Top1), due to its known role in maintaining chromatin stability, we performed rescue experiments to determine how Top1 re-expression in the early stages of aging, or overexpression in log-phase yeast, influences RLS and rDNA silencing, respectively. Here we demonstrate the importance of Top1 homeostasis in maintaining RENT complex enrichment at the rDNA, and report both structural and enzymatic roles for Top1 in establishing rDNA silencing.

## RESULTS

### Nuclear protein levels change significantly in the early stages of replicative aging

Changes in the intracellular protein levels of very old yeast cells (>25 generations) are influenced by a decrease in global translational efficiency (26), but the changes that occur in the early stages of replicative aging are not yet known. To better understand how the nuclear protein landscape changes during the initial stages of aging, when cells are not yet senescing, we performed a proteomic screen comparing isolated nuclei from young (∼0-2 generations) versus moderately aged (∼6-7 generations) yeast cells (diploid lab strain BY4743) that were isolated from mini-chemostat aging devices (see materials and methods). Following isolation of nuclei, we verified nuclear enrichment by DAPI staining under a fluorescence microscope (Figure S1A) and by confirming high levels of histone acetyltransferase (HAT) activity relative to purified HAT1 as a positive control (Figure S1B). We then performed TMT-mass spectrometry on the purified nuclei and calculated the log-fold change of protein levels in the old nuclei compared to the young nuclei (Figure 1A). Roughly 70% of the ∼1000 proteins identified in the screen were unchanged in the aged samples, while 159 were significantly upregulated and 248 significantly downregulated in the aged samples (Figure 1A, Supplementary File 1). To verify results from the proteomic screen, we first chose one of the significantly upregulated proteins, Hsp104, to tag at the C-terminus with the 13xMyc epitope. Hsp104 was chosen because foci of this chaperone protein accumulate in aged cells as it interacts with damaged and misfolded proteins (33). We aged the *HSP104*-13xMyc tagged strain (MD188) for 36 hr. (∼8 generations) on the mini-chemostat system and performed western blots on whole cell protein extracts (Figure 1B). Comparison of aged cell Hsp104-Myc protein expression to young cell controls showed a significant increase in total Hsp104 expression, although the full-length protein remained unchanged. This result suggested that the 13xMyc epitope tag was likely being processed in the older cells at the whole cell level as Hsp104 protein expression increases and foci accumulate (Figure 1B). Next, GO term analysis was performed for the significantly depleted (Figure 1C) or significantly increased (Figure 1D) proteins using the molecular function and biological pathway GO term activity models from GProfiler (34). Among the significantly decreased proteins that were purified with the nuclei, many were involved in ribosome biogenesis, such as RNA Polymerase I and III subunits and the rRNA processing machinery. We also note significant reduction of ribosomal subunits, which are assembled in the nucleolus. Also of particular interest was downregulation of proteins with roles in the maintenance of chromatin structure and stability, including the core histones and histone variants. Histones are reduced in the later stages of aging (35), and we have now confirmed this with earlier stages of aging, thus further validating results from the proteomic screen. Proteins that were significantly increased with age include many involved in the maintenance of cellular homeostasis, including metabolic processes such as carbohydrate, glucan, NADH, and ethanol metabolism (Figure 1D).

**Figure 1.**
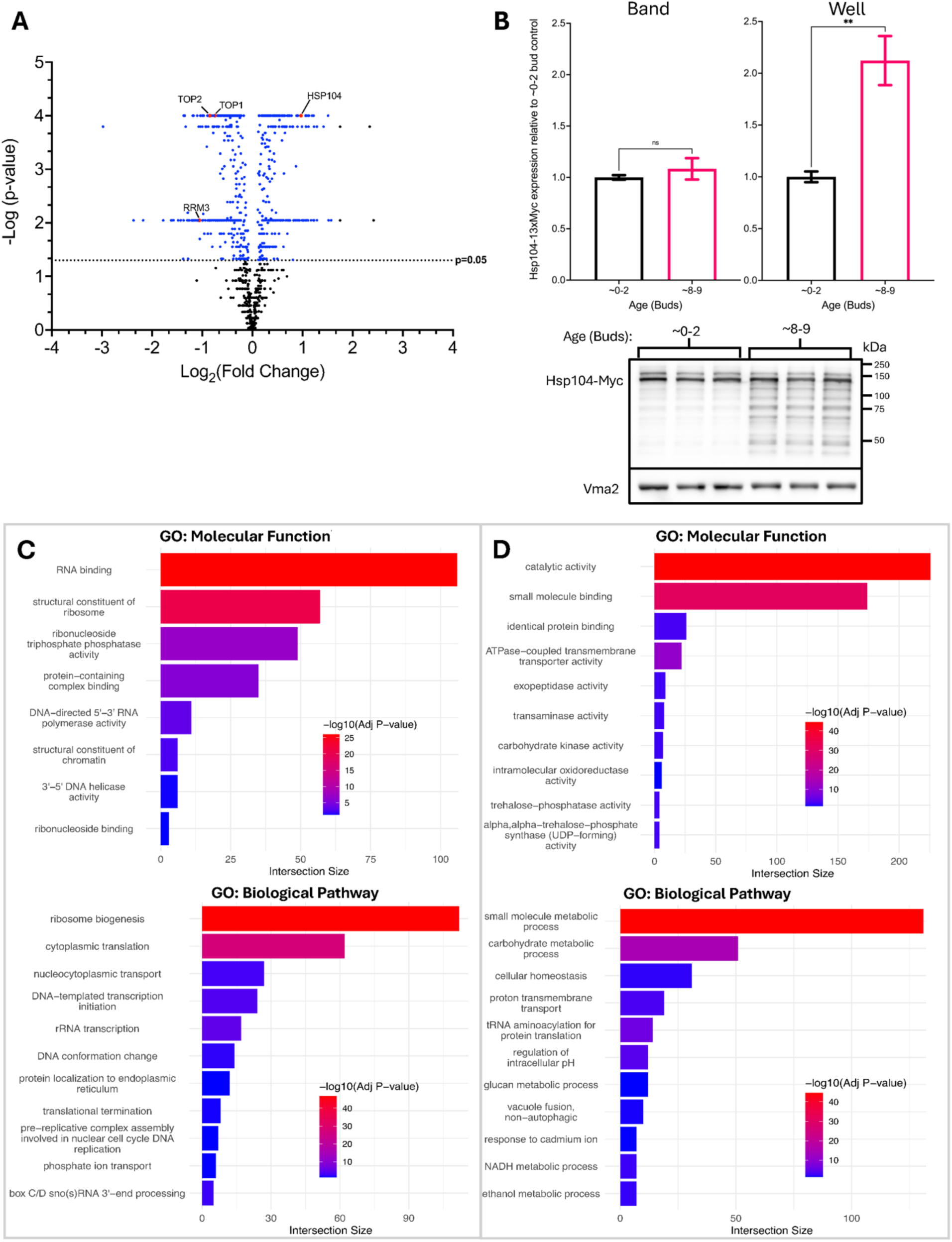
Nuclear protein levels change in early replicative aging. **(A)** -Log fold change in protein expression for nuclei isolated from moderately aged (∼6-7 buds) yeast (BY4743) relative to young (∼0-2 buds) controls obtained using TMT-Mass spectrometry (n=5, -log10 p-values comparing change between old and young cells were calculated for each protein by ANOVA with corrections for multiple comparisons). **(B)** Hsp104-13xMyc protein expression in whole cell extracts from young and aged (∼8-9 generations) MD188 cells (two-tailed student’s t-tests, p=0.2460 and **p=0.0013, n=3). The loading control Vma2 does not significantly change during replicative aging (78). **(C)** GO term enrichment for age-depleted proteins using the molecular function (top) or biological pathway (bottom) models using G-profiler (34). **(D)** Significant GO term enrichment for upregulated proteins.

### Topoisomerase I depletion in early aging is not from decreased transcription or increased protein turnover

Previous work from our lab and others demonstrated the age-associated depletion and redistribution of important rDNA stabilizing proteins such as Sir2 and the Mcd1 subunit of cohesin (30, 31, 36), both of which can extend RLS upon overexpression (30). We thus followed up on age-depleted proteins involved in chromatin stability, as we predicted the depletion of these proteins in early aging would negatively impact RLS. We focused on topoisomerase I (Top1), as it has important dual roles in i) relieving supercoiling associated with DNA replication and transcription genome-wide, and ii) recruitment of Sir2 to the rDNA locus (37–39). We first verified by western blotting that Top1-Myc (strain LP128) was significantly depleted in mother cells replicatively aged for either ∼6 (18 hrs) or ∼9 (36 hrs) generations in the mini-chemostats (Figure 2A). Top1-myc was already strongly depleted to ∼20% within 6 generations, thus validating the proteomic result and also confirming that the 13x-myc tag was not artificially stabilizing the fusion protein during aging.

**Figure 2.**
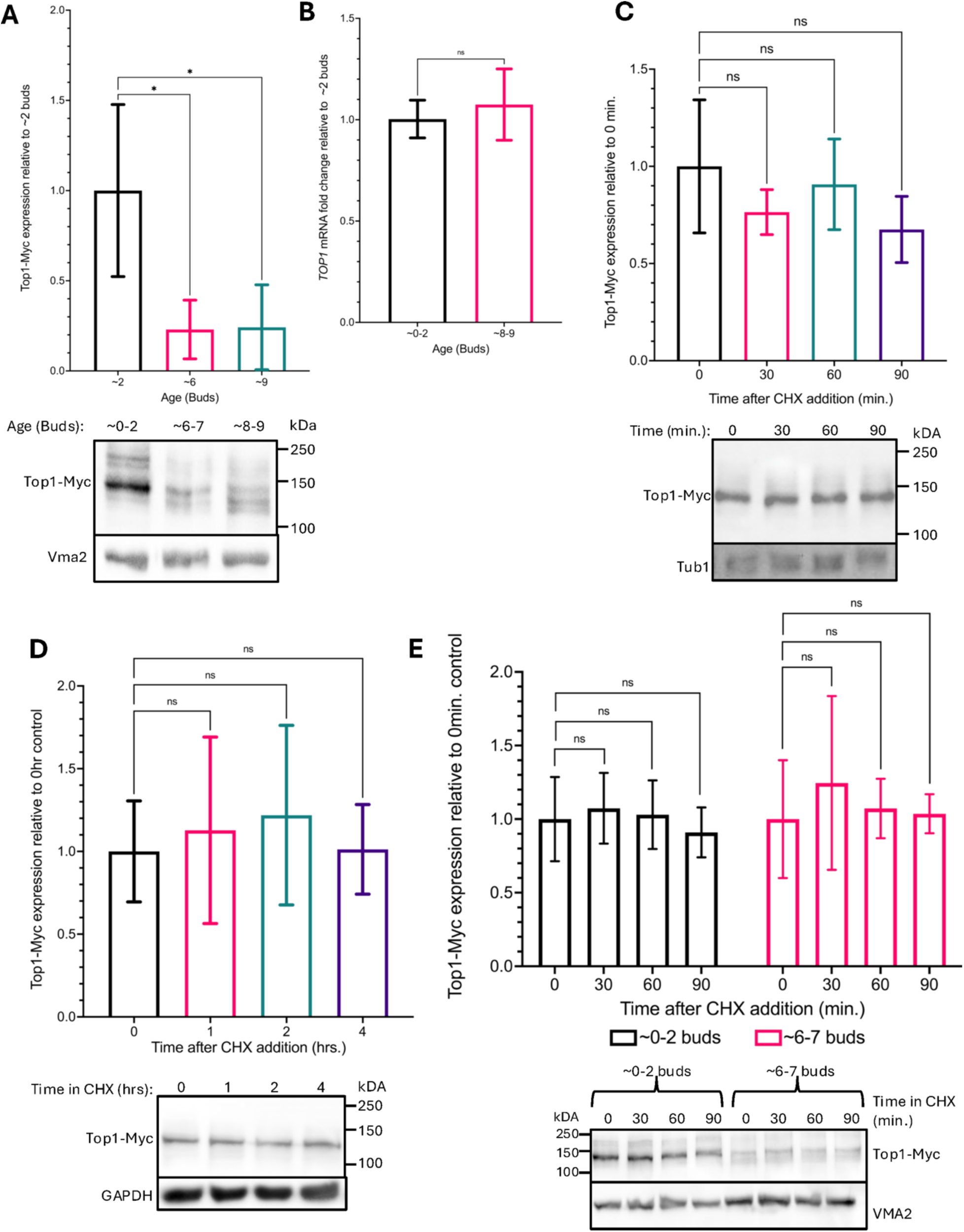
Top1 is depleted at the protein level during replicative aging. **(A)** Quantitation of Top1-13xMyc depletion from LP128 cells aged to ∼6 or 9 generations compared to young controls (∼0-2 generations). (One-way ANOVA with Dunnett’s test for multiple comparisons, *p=0.0454 and *p=0.0482, n=3) **(B)** qRT-PCR results showing mRNA fold change of *TOP1* after ∼8-9 generations relative to their young control (∼0-2 generations) (two-tailed t-test, p=0.4988 n=4). **(C)** Top1-Myc expression after 0, 30, 60, and 90 minutes chase in 250µg/mL CHX relative to the 0 minutes control **(D)** Top1-Myc expression after 0, 1, 2, and 4 hour chase in 250µg/mL CHX relative to the 0 hours control **(E)** Top1-Myc expression in young (∼0-2 buds) and moderately aged (∼6-7 buds) after 0,30,60, and 90 minutes incubation in 250ug/mL CHX relative to the 0 minutes control. All CHX assays were analyzed by one-way ANOVAs with Dunnett’s test for multiple comparisons, adjusted p-values were all ns (p>0.05), n=3. Exact p-values are listed in supplemental Table S4.

We next verified that the 13xMyc epitope tag was not affecting the rDNA silencing function of Top1. Since deletion of *top1* causes a silencing defect in yeast due to a loss of Sir2 from the rDNA locus (15, 39), we performed qRT-PCR for the IGS1 lncRNA that is transcribed from the rDNA when silencing is defective (Figure S2). Relative to an untagged WT control BY4741, the IGS1 lncRNA expression level from the Top1-Myc tagged strain was not significantly changed, while IGS1 lncRNA expression from a *sir2Δ* control strain was significantly increased relative to the wildtype thus demonstrating that the Myc-tag epitope does not significantly impact Top1 rDNA silencing activity.

We next asked if the age-dependent Top1 protein depletion was due to downregulated transcription of the *TOP1* gene, so we isolated RNA from young and moderately aged cells for RT-qPCR assays. *TOP1* mRNA expression levels were unchanged in aged cells (Figure 2B), indicating that decreased transcription or mRNA stability was not the cause of the age induced protein depletion. We also hypothesized that there could be faster Top1 protein turnover induced by aging. To test this idea, a series of cycloheximide (CHX) chase assays was performed. We first verified that a final concentration of 250 µg/ml CHX, commonly used for such experiments in yeast, was sufficient to arrest cell growth in order to confirm the compound was effectively inhibiting translation (Figure S3A). As positive control, a 4-hour CHX assay was performed on a temperature sensitive Cdc13-13xMyc protein with a short half-life of less than 1 hour (Figure S3B) (40). The half-life of Top1 has not been published, so we initially ran a CHX chase experiment on log-phase cells and did not observe any significant turnover of Top1-Myc protein after 90 minutes of CHX incubation (Figure 2C). We therefore increased the maximum incubation time of the cells in CHX to 4 hours and again did not observe any significant changes in Top1-Myc levels (Figure 2D), indicating that Top1 is a very stable protein in young exponentially growing cells. To determine if Top1-myc was destabilized in the early stages of aging, when the protein level was declining, we performed a CHX chase comparison between young and moderately aged yeast cells (∼6 generations). Top1-myc was again depleted in the aged cells, but there was still no significant turnover after 90 minutes in CHX (Figure 2E). Based on these results, the depletion of Top1 during early stages of replicative aging is most likely due to reduced translation.

### Rescue of age-dependent Top1 depletion decreases RLS

Based on previous work showing that overexpression of either Mcd1 (a subunit of the cohesin complex) or Sir2 (a subunit of the RENT and SIR complexes) was sufficient to extend RLS (20, 30), we sought to test whether rescuing the age-dependent depletion of Top1 would also extend RLS. Top1 is important for recruiting Sir2 to the rDNA locus and deletion of Top1 causes silencing defects at the rDNA (15, 39), so we predicted that rescuing Top1 depletion would enhance silencing at the rDNA and extend RLS. Using a haploid strain from the YETI (Yeast Estradiol strain with Titratable Expression) collection that expresses Top1 tagged with 13xMyc (41), we observed ∼2-fold increased protein expression when induced with 2.5 or 5 nM β-estradiol as compared to the endogenous Top1-13xMyc level in a control (border) strain (Figure S4). These strains were compared for RLS with a manual dissection assay on YPD agar containing 10 nM β-estradiol, and surprisingly, RLS of the *TOP1* overexpression strain was significantly lower than the border control strain (Figure 3A). We hypothesized that the elevated Top1 expression could be detrimental due to DNA damage, which was described earlier for a highly expressed copper-inducible *TOP1* system (42). We therefore shifted to expressing *TOP1* expression from its own promoter on a low copy CEN/ARS *URA3* vector. Cells were grown in SC-Ura liquid media and RLS was tracked using an automated microfluidics imaging system, followed by manual counting of the cell divisions. This analysis still revealed a trend toward reduced RLS for WT-*TOP1* vector that was not statistically significant when compared to the empty vector (Figure 3B). We therefore predicted that expressing a catalytically dead Top1-Y727F mutant (Top1-CD), which binds to DNA without forming cleavage-complexes, would help recruit Sir2 to the rDNA without inducing genome-wide damage, thus extending RLS (39). However, the same CEN/ARS plasmid expressing the Top1-CD mutant significantly decreased RLS compared to the empty vector control (Figure 3C). We next considered the possibility that rescuing Top1 expression in aging cells did not extend RLS because Sir2 was also strongly depleted during aging, thus preventing the ability of the Top1 WT or catalytically dead proteins to enhance rDNA silencing and RLS. To address this idea, we enhanced the Sir2 pool by integrating an extra copy of the *SIR2* gene at the *leu2Δ1* locus and then ran the microfluidics experiment again with the Top1-WT and Top1-CD *CEN/ARS* vectors. To our continued surprise, there was a significant decrease in RLS for both *TOP1* overexpression strains compared to the empty vector control (Figure 3D and E), as quantified by a custom-built automated image analysis pipeline. The longer overall RLS observed for the survival curves in panel E compared to panels B and C is due to the increased *SIR2* expression, which strongly extends lifespan (20). Top1 overexpression therefore partially counteracts the RLS extension induced by increased *SIR2* expression.

**Figure 3.**
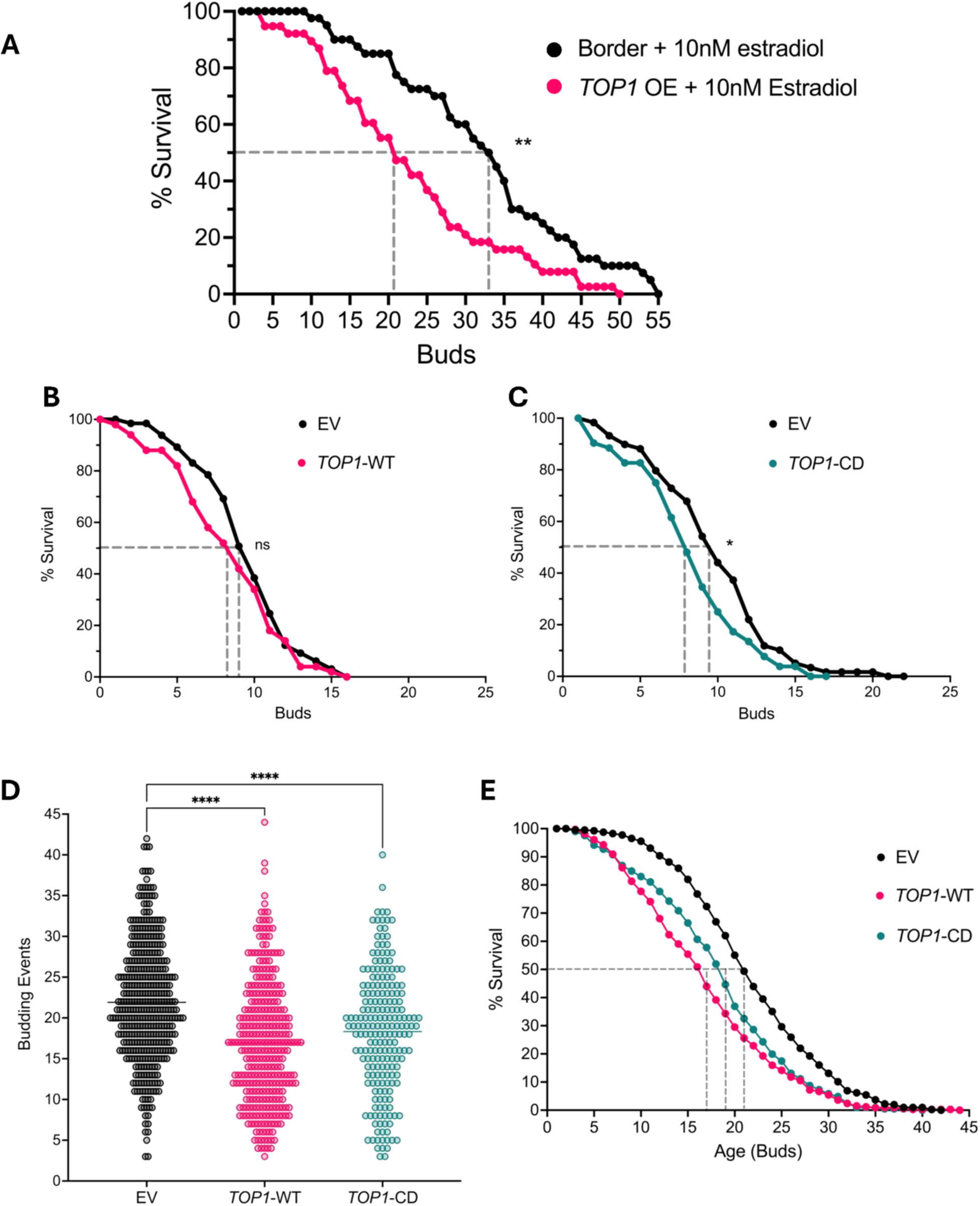
Top1 overexpression to prevent age-associated depletion does not extend RLS. **(A**) RLS measured by manual microdissection for a WT border strain (LP55, n=41) and a Top1 YETI overexpression strain (LP56, n=38) on SC + 10nM β-Estradiol agar plates. Mean RLS= 32.44 (LP55) and 22.79 (LP56), Log-rank test p=0.0012. **(B)** RLS manually counted from microfluidics chips with strains expressing either a low copy empty vector (LP181, n=65) or low copy vector expressing WT Top1 from its native promoter (LP183, n=50). Mean RLS = 9.55 (LP181) and 8.46 (LP183), p=0.2519. **(C)** RLS manually counted from microfluidics chips with strains expressing the vector (LP181, n=59) or catalytically dead (Top1-CD) mutant (LP185, n=52). Mean RLS = 9.85 (LP181) and 8.35 (LP185), p=0.0354. **(D)** Automated counting of budding events for mother cells with an extra integrated copy of *SIR2* and harboring either a low copy pRS416 empty vector (LP257, n=405), a pRS416-WT-Top1 vector (LP263, n=332), or pRS416-Top1-CD vector (LP269, n=206). Mean RLS = 21.9 (EV), 16.9 (Top1-WT) and 18.3 (Top1-CD), Kruskal Wallis test with Dunn’s test for pairwise comparisons with BH corrections for multiple hypotheses testing p<0.0001. **(E)** Survival curve depiction of the automated budding counts for the same strains. Kruskal Wallis rank-sum test p<0.0001.

### Overexpression of a catalytically dead Top1 mutant decreases silencing of an rDNA-flanking reporter gene

Due to the negative effects of Top1 overexpression (both WT and catalytically dead) on RLS, we asked whether this overexpression affected rDNA silencing, as rDNA silencing is closely linked to RLS (18). To test the effects of Top1 overexpression on rDNA silencing, we performed silencing assays on strains carrying a modified *URA3* reporter gene (*mURA3*) integrated into unique chrXII sequence adjacent to the leftmost rDNA repeat (strain YNM44) (Figure 4A) (43). Repressive chromatin spreads from the Fob1 binding sites (TER1 and TER2) into the adjacent *mURA3* reporter. In this assay, strains with weakened silencing of the *mURA3* reporter grow faster on SC media lacking uracil (SC-Leu-Ura) but slower on media that contains 5-fluoroorotic-acid (SC-Leu+FOA). When we transformed YNM44 with either a low copy CEN/ARS or a high copy 2μ *LEU2* plasmid expressing WT *TOP1*, there was a slight weakening of silencing that became more severe with a 2μ Top1-CD catalytic mutant plasmid. This was indicated by increased growth on SC-Leu-Ura plates and weakened growth on SC-Leu+FOA plates (Figure 4A). CEN/ARS and 2μ vectors expressing *NET1* or *SIR2* acted as negative and positive controls, as it has been published by our lab that *NET1* overexpression decreases rDNA silencing, while *SIR2* overexpression enhances rDNA silencing of the *mURA3* reporter (19). As previously discussed, strains with a *TOP1* deletion exhibit an rDNA silencing defect similar to that of *SIR2* deletion strains (15). To determine if expression of Top1-WT or Top1-CD from the plasmids was sufficient to rescue the *top1Δ* silencing defect phenotype, we deleted *TOP1* from YNM44 and transformed with empty vector, Top1-WT or Top1-CD plasmids. The WT Top1 CEN/ARS and 2μ plasmids both restored silencing as compared to the empty vector, indicated by reduced growth on SC-Leu-Ura and improved growth on SC-Leu+FOA (Figure 4B). Interestingly, the catalytically dead Top1-CD did not restore rDNA silencing to the *top1Δ* strain, indicating that the catalytic activity of Top1 is important for establishing silencing. Previous studies reported that the catalytic activity of Top1 was not required for Sir2 recruitment to the rDNA, which raises the question of why the activity would be required in the silencing reporter strains. This question will be addressed in the discussion.

**Figure 4.**
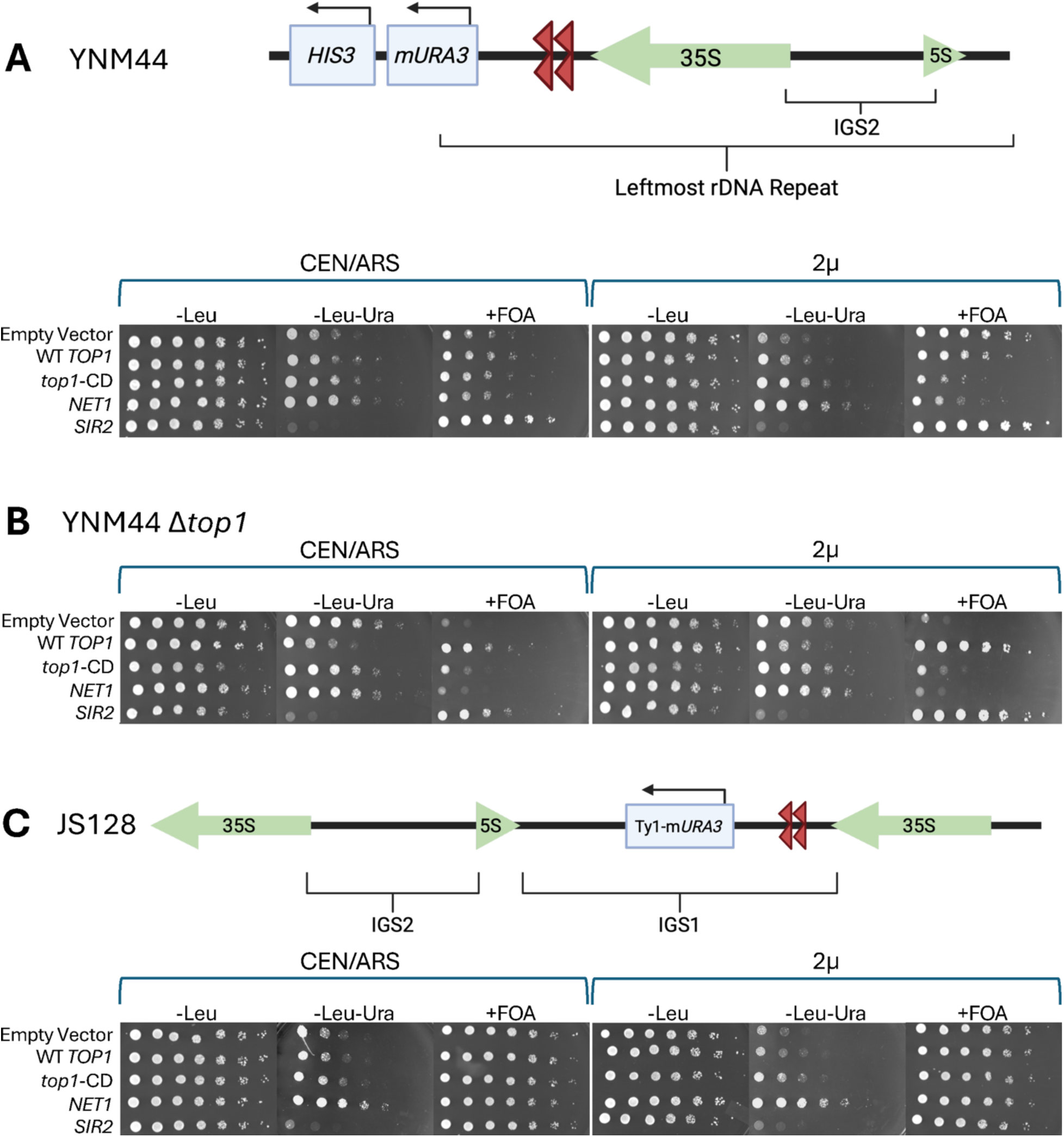
Top1 overexpression negatively affects rDNA silencing. Spot assays on SC-Leu, SC-Leu-Ura and SC-Leu+FOA plates to assess silencing of a *mURA3* reporter gene when *TOP1*, *top1-CD*, *NET1* or *SIR2* are overexpressed from a low copy (CEN/ARS) or a high copy (2μ) *LEU2* vector in the following strains: **(A) YNM44** (*mURA3* reporter is directly <u>flanking the rDNA leftmost repeat)</u>, **(B) YNM44 *top1Δ***, **(C) JS128** (Ty1-*mURA3* reporter integrated at the <u>IGS1 region of an rDNA repeat located with the tandem array)</u>.

Because the overexpression of Top1 had negative effects on silencing of an *mURA3* reporter gene flanking the rDNA, we next asked if *TOP1* overexpression would also weaken silencing with the reporter gene positioned at the IGS1 region within the rDNA array. Previous literature has shown that RNA Pol II binding and ncRNA transcription at IGS1 were elevated when silencing of this region was lost via deletion of *sir2* (14). We therefore hypothesized that the Top1 overexpression vectors should also weaken reporter gene silencing at the internal IGS1 region. We found that when either WT Top1 or Top1-CD was overexpressed, silencing of the *mURA3* reporter within IGS1 was mildly derepressed (Figure 4C). Similar to the left flanking reporter, overexpression of the Top1 catalytic mutant had a stronger effect on IGS1 reporter silencing than the WT Top1.

### Top1 overexpression removes Sir2 from the rDNA

We next asked if the overexpression of Top1 or Top1-CD influenced the binding of Sir2 at the intergenic spacer regions of the rDNA, thereby providing a mechanism for the weakened silencing and shortened RLS. Using ChIP assays, we found that overexpressing the wildtype Top1 or Top1-CD mutant from a low copy CEN/ARS vector was sufficient to significantly decrease the enrichment of Myc-tagged Sir2 at both IGS1 and IGS2, with the catalytically dead mutant having a greater effect (Figure 5A and B). This was consistent with the stronger negative effect of overexpressing the mutant on rDNA silencing.

**Figure 5.**
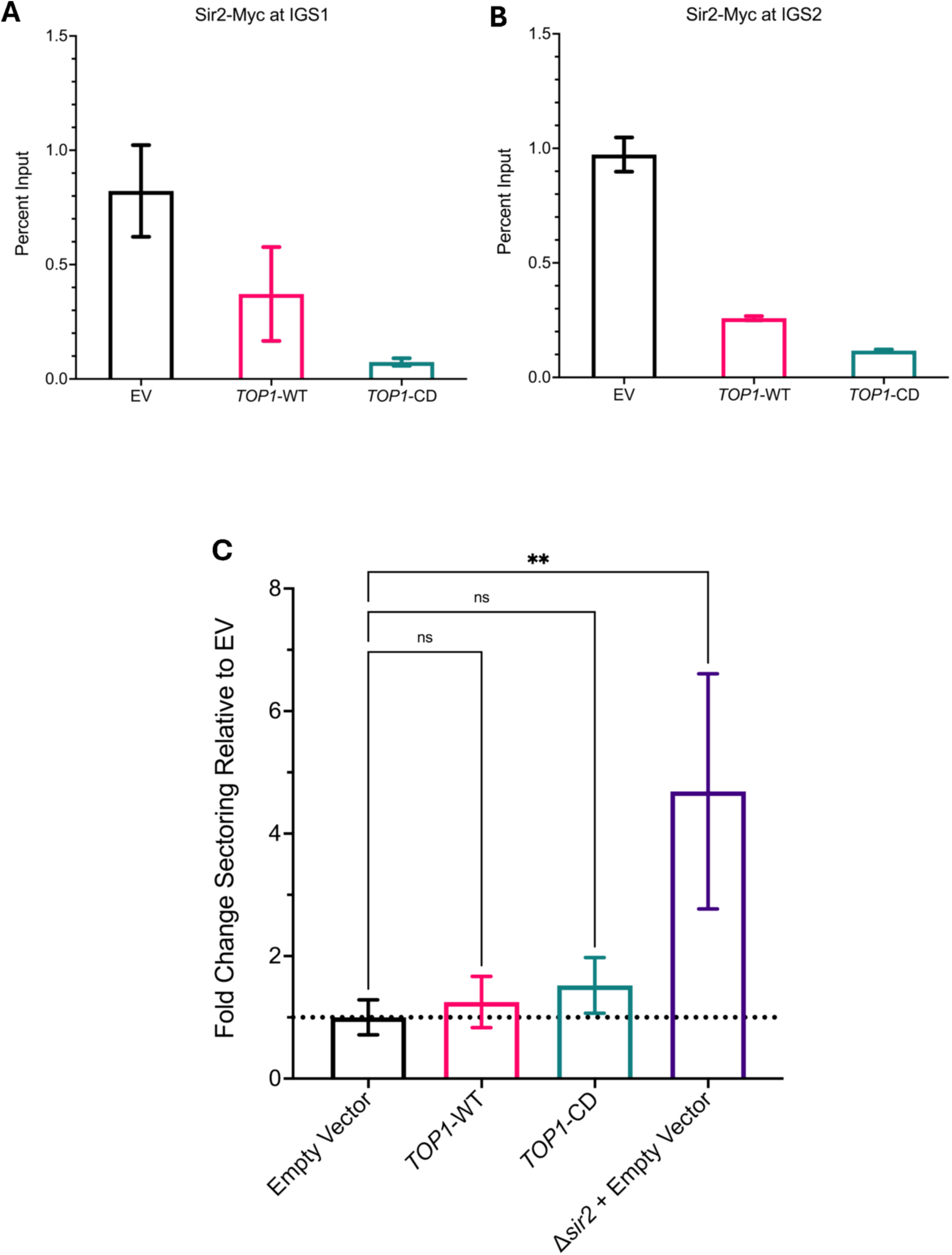
Top1 overexpression reduces Sir2 binding at the rDNA and decreases rDNA stability. **(A)** Quantitative ChIP-assay to assess enrichment of Sir2-Myc at the IGS1 region of rDNA, 3 technical replicates. **(B)** Quantitative ChIP assessing Sir2-Myc enrichment at IGS2. Strain MD209 harbors the empty vector (EV) pRS416, 3 technical replicates. MD208 harbors the Top1-WT vector, and MD210 harbors the Top1-CD vector. **(C)** Frequency of rDNA recombination indicated by loss of an integrated *ADE2* reporter. The frequency of ½ red/white sectored colonies from a strain (AW72) harboring the high copy 2µ empty vector pASC425 was normalized to 1.0. Additional overexpression strains were *TOP1*-WT (AW73) and *top1*-CD (AW74). A s*ir2Δ* deletion strain with the empty vector was used as a positive control for elevated sectoring (AW82). From left to right p=0.9805, p=0.8649, **p=0.0056, one-way ANOVA with Dunnet’s correction for multiple comparisons. Three biological replicates were performed for each strain.

The notable reduction of rDNA silencing along with the disruption of Sir2 recruitment to the rDNA when Top1 was overexpressed led us to ask if there were also negative effects on rDNA stability. To test this, we performed colony sectoring assays with a strain harboring an *ADE2* marker positioned within the rDNA array (20). Loss of this marker from the rDNA due to recombination repair events causes the cells to appear red when plated on media limited for adenine. Colonies that are ½ red and ½ white indicate that the *ADE2* marker was lost upon the first cell division after plating. The high copy WT Top1 and catalytically dead Top1 vectors were transformed into the *ADE2* reporter strain and then tested for marker loss, as compared to the empty vector. While not scored as significant, the *top1* mutant trended toward modestly elevated marker loss, consistent with the weaker rDNA silencing and displaced Sir2 phenotype. As a positive control to ensure we can detect a significant increase in sectoring, we used a s*ir2Δ* strain known to have high rDNA instability (30) (Figure 5C).

Given the role of Top1 catalytic activity in establishing rDNA silencing as well as its ability to dilute Sir2 away from both IGS1 and IGS2, we have proposed a model for the effects of Top1 overexpression at the rDNA locus (Figure 6). In this model, under normal Top1 protein levels, Top1 is bound to Fob1 at the rDNA locus, where Top1 catalytic activity allows the RENT complex to bind and silence RNA Pol II transcription in the intergenic spacer (Figure 6A). However, when *TOP1* is overexpressed, the RENT complex is diluted away from the rDNA array and silencing of RNA Pol II transcription in the rDNA is weakened (Figure 6B), resulting in mild rDNA instability and shortened RLS.

**Figure 6.**
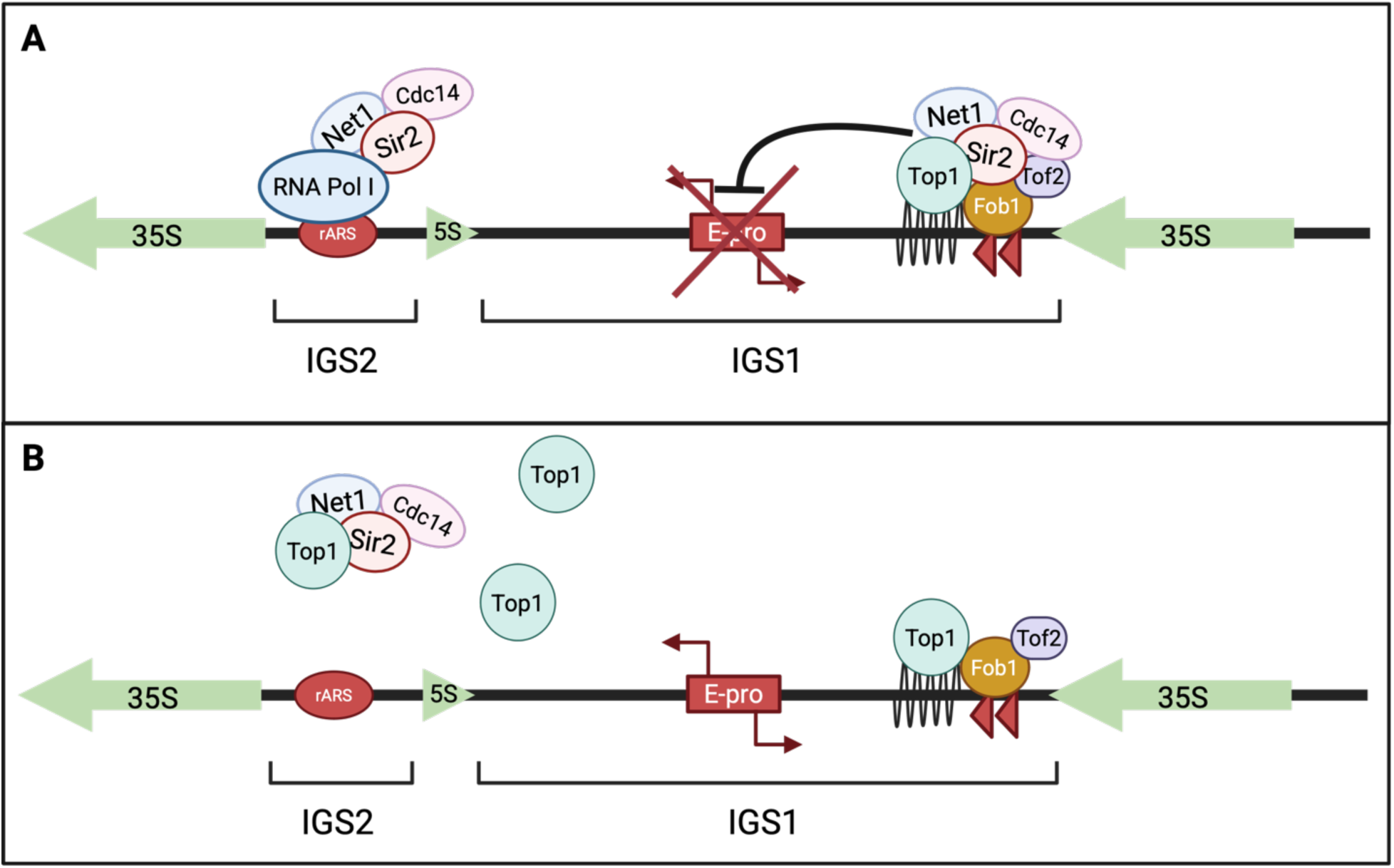
Model for Top1 overexpression disrupting rDNA silencing and RLS. **(A**) Under normal Top1 protein levels, RENT is recruited to IGS1 by interaction with Fob1 and to IGS2 by interaction with RNA Pol I at the rDNA promoter. Top1 catalytic activity enhances RENT enrichment and/or chromatin topology to establish silencing. **(B)** When Top1 is overexpressed, the RENT complex is diluted away from the rDNA array and silencing of RNA Pol II transcription in the rDNA is lost, resulting in expression from E-pro.

## DISCUSSION

### Age-dependent depletion of specific nuclear protein levels: a driver or a consequence of early aging?

While proteomic screens in young and old yeast cells have been reported previously in the literature, our screen was performed on isolated nuclei rather than whole cell extracts and focused on moderately aged cells (∼6-7 generations), giving us novel insight into proteome changes occurring during the earliest stages of replicative aging. A significant portion of the nuclear depleted proteins in early aging were involved in ribosome biogenesis, especially ribosomal RNA processing. This was consistent with previous whole cell aging proteome data showing a significant downregulation of ribosome assembly proteins in cells at ∼20-25 generations (36). Since ribosome biogenesis primarily occurs in the nucleolus, our results with isolated nuclei support the decrease in biogenesis initiating even earlier at ∼6-7 generations. Transcription of the rDNA by RNA Pol I is critical for establishing Sir2-dependent silencing and maintaining stability of the rDNA tandem array (19, 43). The decline in ribosome biogenesis factors in early aging such as RNA Pol I subunits and rRNA processing factors could be directly contributing to the age-associated rDNA instability.

A decline in ribosome biogenesis will also have critical effects on maintaining proteostasis, one of the hallmarks of aging (1). However, the ribosome biogenesis decline is only one aspect of the loss of proteostasis in replicative aging. Proper processing of accumulated protein aggregates is also critical to avoid cell cycle arrest. An increase of protein aggregates in budding yeast decreases RLS but can be rescued by overexpression of chaperone proteins and *CLN3,* a G1 cyclin that is normally present at low levels in very old cells just before they reach senescence (44). Despite the decreases in ribosome biogenesis proteins exhibited in early aging cells, we still detect enrichment of numerous proteins in the aged cells, suggesting that the previously observed general decrease in translation does not apply to all classes of proteins (26). Alternatively, the upregulated proteins could be accumulating due to increased transcription or protein stability. The mechanism of such selective protein enrichment therefore remains an unanswered question in the field. Among the proteins enriched in the aged nuclear preps, we found a significant increase in proteins involved in metabolism, consistent with previous literature that yeast cells will upregulate transcription of genes involved in gluconeogenesis and the glyoxylate cycle (45). While the identification of metabolic proteins accumulated in the aged mother cells was somewhat unsurprising, it is intriguing when we consider that numerous such enzymes have been shown to moonlight as chromatin associated factors that regulate nucleolar activity such as rRNA transcription (46). Validating our screen, one of the most enriched proteins was Hsp104, a stress response protein which has been shown in previous studies to aid in sequestration of damaged proteins to the aging mother cell, and when overexpressed, can partially rescue the RLS of a *SIR2* deletion strain (33).

Additionally, we found that a large set of chromatin stabilizing factors are depleted in the early stages of aging. It is known that specific chromatin stability factors (Mcd1, Sir2, and Sir4) are depleted during aging and that this limited pool of proteins is redistributed from the rDNA to other regions of the genome (30). Through this screen we identified additional chromatin stability proteins, in addition to Top1 discussed in this paper, that were significantly depleted with age such as topoisomerase 2 and Rrm3 helicase, both of which warrant further study. Rrm3 in particular, plays a specific role in unwinding supercoiling at the rDNA replication fork block, where a replication fork and RNA Pol I transcription meet, and is critical for replication fork progression at the rDNA (47, 48). Despite these insights into how the nuclear proteome changes in early aging, it remains unclear if these changes are causal for aging or simply a byproduct of the aging process.

### Mechanisms of Top1 depletion in early replicative aging

The mechanisms behind the observed age-dependent protein depletion remain unknown and the specificity of which proteins become depleted raises intriguing questions for further exploration. Focusing on Top1 depletion specifically, we ruled out decreases in transcription or increased age-induced protein turnover as possible depletion mechanisms, though whether this is true for other depleted proteins needs further investigation. First, we concluded that during the early stages of replicative aging, Top1 mRNA transcript levels are not significantly changed, so transcriptional regulation of Top1 is unlikely to be the cause of Top1 protein depletion with aging. At the RNA-seq level, a recent study found that much of the altered individual protein abundance in old cells correlated with the change in mRNA abundance (49). However, Top1 provides a good example of an exception to this observation showing no change in mRNA abundance with age (Fig. 2B), despite the noted strong protein depletion.

In budding yeast, many proteins are highly stable unless specifically involved in cell cycle regulation (50). Fitting with this data, Top1 is a highly stable protein, even in aged cells. Consistent with the hypothesis that aging-mediated depletion of Top1 is due to a decrease in translation, a previous Ribo-seq study identified a decrease in Top1 translational efficiency during replicative aging (26). However, an independent Ribo-seq study indicated no age-associated change in Top1 (51). The discrepancy between the two ribo-seq findings could be due to differences in the average age of the cells or methods of mother cell isolation. Therefore, the exact mechanism of Top1 translational changes during aging warrant further study. It will also be interesting to test if other age-depleted chromatin related proteins such as Top2 and Rrm3 share the same mechanism of depletion or if they are also specifically impacted by translation efficiency.

A key phenotype of replicatively aging yeast mother cells is slowed cell cycle progression through G1, which is accompanied by progressively enlarged cell volume (52). Artificially slowing down the G1/S transition by deleting the G1 cyclin gene *CLN3* also results in large cells (53). Like our analysis of the nuclear proteome, recent whole proteome SILAC analysis comparing young and old mother cells revealed significant reduction of histones and other chromatin associated factors in the old cells (49). Interestingly, such proteins were also reduced in large *cln3Δ* cells (49), suggesting that at least some of the age-associated protein abundance changes that we observed in nuclei could be related to the enlarged cell size. Interestingly, large *cln3Δ* cells accumulate ERCs as they are passaged for multiple generations, indicating instability of the rDNA array (54). Therefore, even if some chromatin-associated protein depletion in old cells is related to cell size, the consequences on rDNA stability appear to overlap. Given that the cell size scaling of protein abundance is conserved in mammalian cells (49), distinguishing chromatin regulatory processes impacted by cell size or ploidy changes from those specifically impacted by aging will be critical moving forward.

### Less is more: rescuing age-dependent protein depletion does not always extend replicative lifespan

Top1 plays a key role in the maintenance of rDNA stability. In the rDNA locus, Top1 is recruited by Fob1 at IGS1 and enhances recruitment of Sir2 to facilitate rDNA silencing (38, 39). Furthermore, *TOP1* deletion has been shown to produce an rDNA instability phenotype (15, 55–60). We were therefore surprised that rescuing the age-dependent depletion of Top1 did not extend RLS. This decrease in lifespan with overexpression could be due to an increase in DSBs from the formation of Top1 cleavage complexes where Top1 cuts the DNA but does not re-ligate (42, 58). We therefore predicted that the structural role of Top1 in recruiting the RENT complex to the rDNA would be necessary for RLS extension but that too much of the catalytic activity of Top1 would be harmful. Surprisingly, the catalytically dead mutant decreased RLS even more than the wildtype or normal overexpression. Questions therefore remain about Top1 depletion in replicative aging. For example, does this depletion actually benefit the cells as they age because of a requirement to match the levels of key interacting multi-subunit complexes that maintain rDNA stability, including RNA Pol I and the RENT complex (19, 27)? If the stoichiometry between Top1 and these complexes is drastically altered during aging, the rDNA could become more unstable and shorten RLS.

### Top1 plays a role in the establishment of silent chromatin at the rDNA locus through changes in chromatin architecture

To investigate the negative effects of Top1 on RLS, we looked at the impact of Top1 overexpression on an *mURA3* silencing reporter flanking the leftmost repeat of the rDNA, which is positioned adjacent to Fob1 binding sites (TER1/TER2) that recruit the RENT complex (43). The benefit of using this reporter strain is that the *mURA3* reporter is integrated into unique chrXII sequence and not actual rDNA repeat sequence, meaning it is not subjected to the high level of recombinational instability that plagues silencing reporter genes integrated within the rDNA tandem array. *TOP1* overexpression, particularly high copy vectors, weakened silencing at this location. To confirm this effect, we also tested silencing of the *mURA3* reporter integrated at IGS1 within the tandem array, which likely has a different 3D chromatin environment. The weakened silencing effects were the same as observed for the rDNA flanking reporter, indicating that Top1 is likely functioning similarly at both positions.

Consistent with the loss of silencing, overexpression of Top1 decreased Sir2 enrichment at the rDNA intergenic spacers (Figures 5A and 5B). This was consistent with the loss of silencing, but surprising given the scaffolding role for Top1 at the rDNA, which is still present even in the catalytic mutant (39). We therefore hypothesize that Top1 overexpression disrupts stoichiometry of the RENT complex by titrating Sir2 away from the complex (Figure 6). Similarly, in strains where *NET1* has been overexpressed, rDNA silencing decreases, which is likely also from disruption of RENT complex stoichiometry (19). WT Top1 overexpression from a high copy vector only caused a small decrease in rDNA silencing while the catalytic mutant effect was much stronger. This could be explained by the larger effect the catalytic mutant Top1 has on blocking Sir2 from the rDNA that we observed in the ChIP experiments. Another possibility is that overexpression of excess wildtype Top1 is toxic to the cell, so cells are more likely to downregulate the endogenous *TOP1* promoter driving expression from the plasmid, but when expressing a catalytic mutant, they do not have an increase in DNA damage that would trigger such a transcriptional feedback response, leaving more overall Top1 protein to compete Sir2 from the rDNA.

Lastly, we found that overexpression of Top1 from a high copy vector was sufficient to rescue the rDNA silencing defects of an endogenous *TOP1* gene deletion, but the catalytic mutant did not rescue. The same was observed with a *CEN/ARS* set of *TOP1* and *top1-CD* plasmids. This is intriguing, as it indicates that the catalytic activity of Top1 is required for the establishment of rDNA silencing, a novel idea that remains underexplored. Interestingly, RNA Pol I transcription is required for the spreading of rDNA silencing repressive chromatin into the left flanking *mURA3* reporter, even though RNA Pol I transcription does not go past the rDNA genes (19). Additionally, Top1 is required for transcriptional elongation of RNA Pol I through rDNA genes (61). It is therefore possible that supercoiling forced ahead of the transcription machinery, with negative supercoiling accumulating behind RNA Pol I, produces a specialized DNA/chromatin topology downstream of the polymerase that favors RENT enrichment and contributes to the repression of RNA Pol II transcription (62). This hypothesis fits nicely with our data indicating that Top1 catalytic activity is necessary for the establishment of silencing at the left flanking *mURA3* reporter and suggests that the topology of the DNA/chromatin, not just the Sir2 recruitment by Top1 may be important to form the repressive chromatin domain. Supporting this hypothesis, at the silent mating-type loci on chromosome III, Sir2-dependent establishment of silent chromatin is more negatively supercoiled than non-silenced chromatin (63). It makes sense then, that Top1 may be required to relieve the positive supercoiling ahead of the RNA Pol I machinery to establish repressive chromatin downstream that silences transcription by RNA Pol II. Additional work will be needed to elucidate this model for rDNA silencing establishment.

## MATERIALS AND METHODS

### Yeast Strains, Plasmids, and Media

Yeast strains were grown at 30°C in yeast peptone dextrose (YPD) growth medium or synthetic complete (SC) dropout media for plasmid maintenance (64, 65). All strains are listed in Table S1, and all DNA plasmids and primers are listed in Tables S2 and S3 respectively. Proteins for western blotting were tagged with 13xMyc epitopes (EQKLISEEDL) at the C-terminus of the endogenous locus (strains MD188, LP128, MD270, LP280) using PCR and *kanMX4* as a selectable marker for integration of the tag cassette with primers (Table S1). MD188, MD270, and LP280 were constructed using the method outlined in Longtine et al., 1998 (66). For LP128 and LP124, a *TOP1*-13xMyc cassette was PCR amplified from YRH919 genomic DNA, which already harbored the insertion, before transformation into BY4741 and LP56, respectively. Strain LP56 is from the YETI overexpression collection and has an estradiol inducible promoter upstream of the endogenous *TOP1* gene (41). Titratable overexpression of Top1 protein from strain LP124 was confirmed with Western Blots. Tagged strains were verified with both colony PCR and western blots. The high copy 2μ Top1-LEU2 (pAW6) and 2μ Top1-CD-LEU2 (pAW8) vectors were constructed by PCR amplifying the wildtype *TOP1* or catalytic mutant *top1*-CD genes with their native promoters and 3’ termination sequences from plasmids pNK66 and pNK67, respectively (67). PCR products from primers JS4054 and JS4055 were digested with *Asc*I restriction endonuclease and ligated into plasmid pAsc425 (19). Low copy versions of these plasmids were generated (CEN/ARS *TOP1-LEU2* (pAW2) and CEN/ARS *top1-*CD*-LEU2* (pAW4)) by ligating the same PCR products into pAsc415. Plasmids were verified with restriction digests and Sanger sequencing. The *NET1 CEN/ARS LEU2* plasmid pSB794, *SIR2 CEN/ARS LEU2* plasmid pSB764, *NET1 2µ LEU2* plasmid pSB790, and *SIR2 2µ LEU2* plasmid pSB766 were previously described and constructed similarly with PCR and *AscI* ligation into pAsc415 or pAsc425 (19). Strains transformed with plasmids were maintained in selective media to prevent plasmid loss.

### Nuclei Isolation for Proteomic Screen

Approximately 4x10^8^ young or aged cells isolated from the Mini Chemostat Aging Device (MAD; see description below) and nuclei purified using a Yeast Nuclei Isolation Kit (ab206997) from Abcam following the kit procedure with minor modifications. Cells were washed two times with distilled water with centrifugation a 3000 x g at room temperature, then resuspended in 1 ml of Buffer A containing 10 mM DTT and incubated at 30°C for 10 min with gentle shaking. The samples were centrifuged at 1500 x g for 5 min at room temperature and the supernatant discarded. The pellets were resuspended in 1 ml of Buffer B, followed by addition of 10 µl of Lysis Enzyme Cocktail and incubation of 15 min at 30°C for young cells and 18 min for old cells to generate spheroplasts. The samples were then centrifuged at 1500 x g for 5 min and the pellets were kept on ice after discarding the supernatants. The spheroplast pellets were resuspended in 1 ml of Buffer C containing protease inhibitor cocktail from the kit and then homogenized in a Dounce homogenizer. The suspension was mixed by shaking for 30 min at room temperature then centrifuged at 1500 x g for 5 min at 4°C to remove cell debris. Supernatants were collected and then centrifuged at 20000 x g for 30 min at 4° to collect the nuclei, which were resuspended in Buffer C. Purified nuclei were quantified for protein concentration and visualized by DAPI staining under a fluorescence microscope. H3-dependent HAT activity is nuclear specific in yeast, so enzymatic HAT activity from the nuclei was also validated using H3 substrate following the manufacturer’s protocol with provided nuclear extract (NE, 4 mg/mL) as a positive control.

### TMT Mass Spectrometry

Tandem Mass Tag (TMT) labeling of nuclear proteins was performed in the Biomolecular Analysis Facility (BAF) core in the UVA School of Medicine following a standard protocol from the Gygi lab (68), with changes noted here. Protein extracts in 50 mM HEPES pH 8.2, 8 M urea, 50 mM NaCL, 1x protease inhibitor cocktail (Complete Mini, Roche, Basel, Switzerland) were generated from bead beating the nuclei then clarifying by centrifugation at 13,000 rpm for 15 minutes. Protein was quantified with a Qubit 3.0 fluorometer (ThermoScientific). Approximately 100 µg of protein from each sample was reduced with 10 mM DTT at room temperature for 45 minutes, followed by alkylation with 10mM iodoacetamide for 30 minutes in the dark at room temperature. Methanol/chloroform precipitation was performed immediately following alkylation, and the protein pellets were washed 3x with ice-cold methanol and rapidly dried by vacuum centrifugation. The sample pellets were resuspended in 20 µL of 50 mM ammonium bicarbonate solution pH 7.8 by repeated pipetting on ice and digested with an approximate 25:1 sequencing grade trypsin at room temperature for 16 hours. The reaction was stopped by the addition of 2.5 mM trifluoroacetic acid (TFA) to a pH <2 and lyophilized with vacuum centrifugation. The peptides were resuspended in 0.1% TFA and verified at a pH of 3 before peptide desalting with spin tips (Pierce™ C18 Spin Tips & Columns) following manufacturer protocols and again lyophilized. Individual samples were labeled with TMT reagents resuspended in anhydrous acetonitrile at a ratio of 1:4 peptide to TMT reagent. The labeled peptides were individually desalted to remove excess TMT reagent, dried with vacuum centrifugation and combined to a total volume of 40 µl in 0.1% TFA. Mass spectrometry was run on an LC-MS (Thermo Electron Q Exactive HF-Xmass spectrometer) system with an Easy Spray ion source connected to a Thermo 75 μmx 15 cm C18 Easy Spray column. 15 μL of the extract was injected and the peptides were eluted from the column by an acetonitrile/0.1 M formic acid gradient at a flow rate of 0.3 μL/min over 2.0 hours. The nanospray ion source was operated at 1.9 kV. The digest was analyzed using the rapid switching capability of the instrument acquiring a full scan mass spectrum (120K resolution) to determine peptide molecular weights followed by production spectra (30 HCD at 7.5 K resolution, 100ms fill, 35 NCE, 1.4 Da isolation) to determine amino acid sequence in sequential scans. This mode of analysis produces approximately 25,000 MS/MS spectra of ions ranging in abundance over several orders of magnitude. Not all MS/MS spectra are derived from peptides. The data were analyzed by database searching using the Sequest search algorithm in Proteome Discoverer 2.2 against Uniprot yeast (10/23/19). Identifications and TMT reporter quantification were performed in Scaffold. The samples produced identifications for ∼1000 proteins. The log fold change between young and old cells was calculated and adjusted p-values were calculated. Analyzed results of proteomic data is available in supporting information Table S5.

### Isolation of Aged Yeast Cells

A <u>M</u>ini Chemostat <u>A</u>ging <u>D</u>evice (MAD) was set up based on designs and protocols from the Dunham Lab and Calico Labs (32, 69). 300 mL cultures were grown shaking in YPD at 30°C overnight until an OD_600_ of ∼1.2-1.7 was reached (log phase). Cells were pelleted and counted by hemocytometer. For each mini-chemostat tube (each chemostat run has 12 tubes), 250 million log phase yeast cells were washed 3 times with 1x PBS/0.25% PEG-3350. Cells were resuspended in 500 µL of 1xPBS, then 500 µL of PBS with 4 mg/mL biotin (Thermo-Scientific EZ-Link™ Sulfo-NHS-LC-Biotin) was added. Cells were biotinylated for 30 minutes with slow rotation at room temperature and then washed twice with 1xPBS to remove unbound biotin. Cells were transferred to 50 mL of YPD media and incubated while shaking at 30°C for 3 hours to recover. Cells were then pelleted and washed with 1xPBS, then resuspended in 600 µL of 1xPBS. 50 µL of magnetic-streptavidin beads (Invitrogen Dynabeads™ MyOne™ Streptavidin C) suspended in PBS were added to each 600 µL of biotinylated cells and incubated with slow rotation at room temperature for 30 minutes. After bead binding, tubes were placed on a magnetic rack, and cells that did not have any beads bound (unbiotinylated daughter cells) were removed. These unbiotinylated daughter cells were kept as the “young control”. The remaining bead-bound cells were loaded into the chemostat tubes and bound to the sides by strong neodymium magnets. Mother cells were incubated in the chemostat device for the indicated amount of time with fresh SC media continuously added with a Watson Marlow 205S peristaltic pump and aerated by aquarium air pumps. Air flow also keeps the daughter cells from settling at the bottom, so they are washed away into waste containers. After aging the mother cells, the content of each chemostat tube was harvested, and magnetic racks were used to collect the mother cells and wash away remaining daughter cells. The average age of the mother cells was determined using calcofluor white staining (incubated in 1 mg/mL calcofluor for 15 minutes, washed and resuspended in 1xPBS), and imaged at 60x objective with an EVOS M7000 microscope from Invitrogen (Thermo Fisher Scientific).

### Real-time PCR (RTPCR)

Total RNA from ∼1x10^8^ cells was isolated using the hot acid phenol method and resuspended in DEPC treated H_2_O (70). 1 μg of RNA was used for cDNA synthesis with the Verso cDNA Synthesis Kit (ThermoFisher) and cDNA was diluted 3-fold in nuclease free H_2_O. Quantitative RT-PCR to determine *TOP1* transcript levels was performed on an Applied Biosystems StepOnePlus Real Time PCR system with primers JS3398 and JS3399. Log fold change was calculated using the ΔΔC_t_ method, where *TOP1* signal was normalized to *UBC6* signal (primers JS3976 and JS3977), and the normalized *TOP1* from old cells was compared to the average of the young control. RTPCR was performed on 3 biological replicates.

### Western Blots

Total protein extracts from ∼5x10^8^ cells were isolated from thawed cell pellets using a protocol adapted from Dunn and Wobbe (71). Pellets were resuspended in 400 μL of lysis buffer (50 mM HEPES pH 7.6, 10% Glycerol, 10 mM EDTA, 0.5 M NaCl, 1% Triton X-100, 5 mM DTT, 1X protease inhibitor cocktail, 1 mM PMSF) and lysed with an equal volume of acid washed glass beads by vortexing (MPI Fastprep-24 classic bead beating lysis system) for 4x45 seconds with one minute of rest in between (only 3x45 seconds for old cells). Lysates were cleared by centrifugation at 14000 rpm for 10 minutes in an Eppendorf microcentrifuge at 4°C, and supernatants were transferred to fresh microfuge tubes. Protein concentration was measured by Bradford Assay with a UV-vis spectrophotometer (BioMate3, ThermoFisher Scientific). One volume of 5x Laemmli buffer was added to 4 volumes of cleared protein lysate. A volume of lysate containing 20 μg of protein was loaded onto 10% SDS-page gels and run for ∼90 minutes at 120V. Protein was transferred to a PVDF membrane (Millipore Sigma™ Immobilon™-P) by wet transfer in 1x Tris-Glycine buffer at 30V for 16 hours at 4°C in a Biorad mini-gel tank. Membranes were blocked in 5% non-fat dried milk/1xTBST for 1 hour at room temperature. Primary antibodies such as c-Myc Monoclonal Antibody (9E10), Vma2 Monoclonal Antibody (13D11B2, RRID AB_2536202), GAPDH Loading Control Monoclonal Antibody (GA1R, RRID AB_10977387), or alpha Tubulin Monoclonal Antibody (YL1/2, RRID AB_2210201)) were diluted 1:5000 in the milk/1xTBST blocking solution and incubated/rocked with the Immobilon membrane for 1 hour at room temperature. Membranes were next washed 3x5 minutes in 1xTBST then incubated in secondary antibody (Anti-Mouse IgG (H+L), HRP Conjugate, Promega) diluted 1:5000 in 5% skim milk/1xTBST for 45 minutes at room temperature. Membranes were washed again, and protein bands of interest were visualized by chemiluminescence (Immobilon Western Chemiluminescent HRP Substrate) on a Bio-Rad Chemidoc MP system using the automatic exposure settings. Molecular weights were estimated using a protein ladder (Precision Plus Protein Dual Color Standards, Bio-Rad). Images were quantified in ImageJ, where boxes of equal size were used to measure band intensity and background. The background-subtracted 13xMyc bands were normalized to the respective loading control signal (also background corrected). The change in 13xMyc signal for each sample was normalized to the mean of the young cells. Western blots for age-dependent protein changes were performed with 3 biological replicates, and representative images are shown.

### Cycloheximide Chase Assays

Cycloheximide (CHX) chase assays were performed according to a protocol from the Rubenstein lab (72). Timepoints for cycloheximide chase assays were taken at 0, 30, 60, and 90 minutes or 0, 60, 120, and 240 minutes. For cycloheximide chase assays with aged cells, samples were harvested directly from the chemostat and immediately subjected to CHX incubation. After CHX incubation at 250 nM, cells were pelleted and frozen on dry ice. Frozen pellets (∼2.5 OD_600_ units) were used for Western blotting via the protocol previously described. Quantification was performed as previously described with GAPDH or Tub1 loading controls for young cells and Vma2 as the loading control for aged cells. All cycloheximide chase assays were quantified with 3 biological replicates, and representative western blot images are shown.

### Replicative Lifespan Assays

Traditional manual RLS plate assays were performed on agar plates based on a protocol from the Kaeberlein lab (73). ∼50 dividing cells from each strain were arranged in a grid on SC agar plates containing 10 nM β-estradiol (for overexpression with the YETI strains) using a Nikon Eclipse E400 microscope outfitted with a manual micromanipulator and fiber optic dissection needle. After being placed on the agar plates and incubated for ∼1 hour, the mother cells are removed to the side of the plate and the new virgin daughter cells become the cohort of mother cells to be used for the experiment. Microdissections were performed every hour by pulling the smaller daughter cell from the mother cell and moving it to the side of the plate with the microdissection needle. In between generations, plates were incubated at 30°C and then moved to 4°C each night to stop the cell division until continuing in the morning. Mother cells that stopped budding within the first 2 generations were censored to avoid incorporating already dead cells. Once a mother cell ceased dividing for three generations, it was marked as dead. Plates were monitored until all mother cells had stopped dividing.

### Microfluidics Replicative Lifespan Assays

Cells were grown to log phase in SC-Ura media to maintain the *TOP1* expression plasmid or empty control plasmids that harbored a *URA3* selectable marker. Cells were diluted to an OD_600_ of 0.1 in filtered SC-Ura media and loaded into the Y-shaped traps of the commercial microfluidics chips that are embedded on standard size microscope slides (iBiochip Automated Dissection Kit). SC-Ura media was run through the microfluidics chip at a constant flow of 2 μL/minute to provide fresh media to the cells and wash away the new daughter cells. The chips/slides were placed in an onstage incubator set to 30°C. The Y-shaped nature of the traps in the microfluidics chip ensured that the cells trapped were virgin daughter cells (74). Images of each well were taken every 10 minutes with the 40x objective on an EVOS M7000 motorized stage microscope (Invitrogen, Thermo Fisher Scientific). Images were concatenated into time lapse videos using the Image-J import image sequence function and saved as AVI files. For manual counting of budding events from the movies, the AVI file names were blinded, and budding events were counted for ∼60 mother cells per condition. Budding events were only counted for mother cells that entered the trap within the first 12 hours of the experiment to avoid counting buds for the daughter cells derived from old mothers. Traps where multiple cells had entered were also discarded to avoid incorrect counting of budding events. For both the plate assays and the manually counted microfluidics videos, Kaplan-Myer survival curves were generated using Oasis2 (75). Significant differences in lifespan were calculated using a log rank test. For microfluidics experiments that were analyzed with automated counting, we used a custom-built program described below. Significant differences in lifespan were calculated in MicroBREW using a Kruskal Wallis test to compare three or more independent groups without assuming normality and followed the test with Dunn’s t-test for pairwise comparisons with Bonferoni-corrections for multiple hypotheses testing.

To quantify replicative lifespan (RLS) using a high-content imaging platform, we developed a custom image processing pipeline in MATLAB. The analysis was performed in two main steps. In the first step, each image frame was processed to detect and record the positions of individual microfluidic traps. Briefly, edge detection (“*edge*”, default parameters) was applied to identify trap boundaries, and the resulting images were converted to binary format. Morphological operations, including dilation and hole filling (“*imfill*”, default settings), were used to refine object structures. Labeled binary images were then generated using “*bwlabel*” to assign unique identifiers to each detected object. A size threshold filter was applied to exclude large artifacts, such as those arising from chains of budded cells. For each identified trap, spatial coordinates (x, y) were extracted and used to track the trap location consistently across all frames in the time-lapse movie.

In the second step, four regions of interest (ROIs) were defined for each trap based on the spatial coordinates. These included one central circular region to track the entry and position of the mother cell, and three adjacent kite-shaped regions to capture signals associated with budding events. Within each of the four ROIs, background was corrected by subtracting the mode of the signal intensity distribution, assigning zeros if the values resulted in negative. The background-corrected pixel intensities were then aggregated and mapped to the corresponding trap ID and frame number for downstream analysis.

Signals generated in MATLAB were further processed in R to identify mother cell entry, cell death, and budding events for each trap. For each trap, the signal from the central ROI was first smoothed using a simple moving average and normalized to the pixel intensity in the first frame of the time series. Peaks in the normalized signal were detected using the ‘*findpeaks’* function with a stringent threshold to reduce false positives due to noise. The first identified peak marked the time of mother cell entry, and the corresponding frame ID was recorded. To detect budding events, the signals from the three kite-shaped ROIs were smoothed (*post-entry*) using a moving average, and peaks were identified using ‘*findpeaks’*, with a minimum peak distance of six frames to reflect the expected time between budding events. Mother cell death was inferred from the central ROI by calculating the first-order lag difference; a sustained drop in the signal was used to indicate cell death. For each trap, the number of budding events detected between the mother cell entry and death was recorded per ROI. Additionally, the start and end points of each peak were captured to eliminate overlapping events, and peak heights were recorded to estimate the magnitude (size) of each budding event.

### Silencing Assays

Cells were patched onto SC-Leu agar plates and incubated overnight at 30°C for ∼24 hours. Cells were scraped up from the agar surface and resuspended in 1 mL of sterile H_2_O in microfuge tubes. The OD_600_ was measured with a UV-Vis spectrophotometer (BioMate3, ThermoFisher Scientific), and each sample was subsequently diluted and normalized to an OD_600_ of 1. A series of 1:5 dilutions of cells in sterile water was made for a total of 7 dilutions. 5 µL of each dilution was then spotted in rows onto SC-Leu, SC-Leu-Ura, or SC-Leu+FOA (5-fluoroorotic acid) agar plates to track the level of silencing of a modified *URA3* reporter gene (*mURA3*) that was integrated either adjacent to the rDNA locus or within the IGS1 intergenic spacer (16, 43). Plates were incubated for 72 hours at 30°C. Images to visualize silencing effects were taken with a Fluorochem-Q imager (Protein Simple).

### Chromatin Immunoprecipitation (ChIP) assays

ChIP was performed based on a standard protocol used in the Smith lab (76). Briefly, log phase yeast cells were crosslinked with 1% formaldehyde for 20 minutes, pelleted, washed, and resuspended in lysis buffer (50 mM HEPES, 140 mM NaCl, 1% Triton X-100, 1 mM EDTA, 0.1% SDS, 0.1 mM PMSF, and 1X protease inhibitor cocktail; Sigma [Sigma Chemical], St. Louis, MO). The cell mixtures were then disrupted with glass beads, followed by sonication in a Diagenode Bioruptor for 15 minutes with 30 seconds on / 30 seconds off cycles, followed by centrifugation at full speed in an Eppendorf microcentrifuge to collect the soluble supernatants. The protein content of the cell lysate was measured using Pierce Bradford Plus protein assay reagent (Thermo Scientific #1856209) and 1 mg of the chromatin lysate was used for each ChIP sample. For Myc-tagged samples, 5 μl of 9E10 anti-Myc antibody (EMD Millipore, catalog #05-419) was used for immunoprecipitation overnight at 4°C, and 1/10^th^ of the supernatant volume was kept as input. The next day, immunoprecipitated lysates were incubated with 25 μl protein G magnetic beads (Thermo Scientific #88848) for 2 hrs. at 4°C followed by washing. The DNA was then eluted in elution buffer and reverse-crosslinked overnight at 65°C. DNA samples were purified using PureLink PCR spin columns (Invitrogen, Carlsbad CA). ChIP DNA was quantified using real-time PCR (Step-One Plus Applied Biosystem) and normalized to the input DNA PCR signal. The percentage (Input %) value for each sample is calculated using % of input = 100*2^(adjusted input − Ct(sample))^. 3 technical replicates of ChIP assays were performed with 1 biological replicate.

### Sectoring Assays

Cells were grown to log phase overnight in SC-Leu liquid media. Overnight cultures were pelleted and then washed with sterile water. Cells were then counted by hemocytometer and diluted such that when 100 µL of the diluted cells were plated onto SC-Leu with a limiting concentration of 80 µM adenine, yielding ∼400 colonies/plate. 3 biological replicates from different colonies were spread onto 50 plates for each strain. The agar plates were incubated at 30°C for 72 hours, then 4°C for 24 hours to fully develop the red coloring of the Ade^-^ cells. ½ sectors (half white/half red colonies) and ¼ sectors (¼ red/ ¾ white colonies) indicating a loss of *ADE2* after the first or second cell division were counted using an Olympus SZ-PT microscope. The total number of colonies per plate was automatically counted using OrganoSeg (77). Entirely red colonies were excluded from total colony counts as they had lost the *ADE2* marker before the first cell division. The percent sectoring frequency for each plate was calculated using the equation below. The fold change in sectoring frequency was calculated relative to the mean of the empty vector control. The sectoring assay was performed on 3 biological replicates in 3independent experiments.

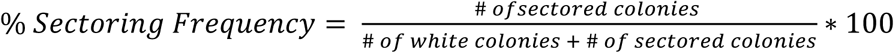

### Statistical Analyses

Statistical analysis for the proteomic screen was performed using a one-way ANOVA with corrections for multiple comparisons to find significant differences (p<0.05) in protein levels of moderately aged cells compared to old cells. 5 biological replicates of aged cell populations were compared to 5 biological replicates of young cell populations. For RT-qPCR and Western blot comparisons of relative mRNA fold change or protein abundance, we performed t-tests (two groups) or one-way ANOVAs (>2 groups) with Dunnett’s test for multiple comparisons. We normalized protein or mRNA abundance to the average of the control group for each value and compared the treatment groups to the control group for each statistical analysis. For western blots and qPCR, we used 3 biological replicates per treatment group. For the sectoring assay, we normalized the sectoring frequency of each sample to the mean sectoring frequency of the empty vector control group and compared experimental groups to the control group using one-way ANOVA with corrections Dunnett’s test for multiple comparisons. For comparison of RLS survival curves we performed log-rank tests. Comparison of mean RLS was performed by Kruskal-Wallis test with Dunn’s test for pairwise comparisons and with BH corrections for multiple hypotheses testing. Graphical representations of p-values are pictured as follows: (ns = p≥0.05, * = p<0.05, ** = p<0.01, *** = p<0.001, **** = p<0.0001).

## Supporting Information

This article contains supporting information.

## Acknowledgments

We thank Ignacio Gutiérrez for advice on setup of the microfluidics system, Maitreya Dunham and Joe Armstrong for advice on setup of the mini-chemostat system. We also thank Sandra Bahr and Griffin Kim for distributing the YETI strains and Dan Gottschling and Nyun Kim for providing additional yeast strains and plasmids. We thank Ani Manichaikul for her feedback on statistical analyses. L.N.P. was supported by NIH predoctoral fellowship F31AG081044. C.T.L. and A.E.W. were supported in part by Cell and Molecular Biology (CMB) NIH training grant T32GM008136. K.K. was supported by NSF grant 1950374. The study was also supported by NIH grant R01GM075240 to J.S.S. and U54CA274499 to K.A.J. This work also utilized proteomics service in the Biomolecular Analysis Core Facility which is supported by the University of Virginia School of Medicine, Research Resource Identifies (RRID):SCR_025476, with co-pay support from the UVA Comprehensive Cancer Center Support grant P30CA044579.

## REFERENCES

1. López-Otín, C., Blasco, M. A., Partridge, L., Serrano, M., and Kroemer, G. (2023) Hallmarks of aging: An expanding universe. Cell. 186, 243–278

2. Mortimer, R. K., and Johnston, J. R. (1959) Life Span of Individual Yeast Cells. Nature. 183, 1751–1752

3. Longo, V. D., Shadel, G. S., Kaeberlein, M., and Kennedy, B. (2012) Replicative and chronological aging in Saccharomyces cerevisiae. Cell Metab. 16, 18–31

4. Smith, J. S., Brachmann, C. B., Celic, I., Kenna, M. A., Muhammad, S., Starai, V. J., Avalos, J. L., Escalante-Semerena, J. C., Grubmeyer, C., Wolberger, C., and Boeke, J. D. (2000) A phylogenetically conserved NAD+-dependent protein deacetylase activity in the Sir2 protein family. Proceedings of the National Academy of Sciences. 97, 6658–6663

5. Landry, J., Sutton, A., Tafrov, S. T., Heller, R. C., Stebbins, J., Pillus, L., and Sternglanz, R. (2000) The silencing protein SIR2 and its homologs are NAD-dependent protein deacetylases. Proc Natl Acad Sci U S A. 97, 5807–5811

6. Imai, S., Armstrong, C. M., Kaeberlein, M., and Guarente, L. (2000) Transcriptional silencing and longevity protein Sir2 is an NAD-dependent histone deacetylase. Nature. 403, 795–800

7. Ghidelli, S., Donze, D., Dhillon, N., and Kamakaka, R. T. (2001) Sir2p exists in two nucleosome-binding complexes with distinct deacetylase activities. EMBO J. 20, 4522–4535

8. Tanny, J. C., Kirkpatrick, D. S., Gerber, S. A., Gygi, S. P., and Moazed, D. (2004) Budding yeast silencing complexes and regulation of Sir2 activity by protein-protein interactions. Mol Cell Biol. 24, 6931–6946

9. Moazed, D., Kistler, A., Axelrod, A., Rine, J., and Johnson, A. D. (1997) Silent information regulator protein complexes in Saccharomyces cerevisiae: a SIR2/SIR4 complex and evidence for a regulatory domain in SIR4 that inhibits its interaction with SIR3. Proc Natl Acad Sci U S A. 94, 2186–2191

10. Moretti, P., Freeman, K., Coodly, L., and Shore, D. (1994) Evidence that a complex of SIR proteins interacts with the silencer and telomere-binding protein RAP1. Genes Dev. 8, 2257–2269

11. Gartenberg, M. R., and Smith, J. S. (2016) The Nuts and Bolts of Transcriptionally Silent Chromatin in *Saccharomyces cerevisiae*. Genetics. 203, 1563–1599

12. Shou, W., Seol, J. H., Shevchenko, A., Baskerville, C., Moazed, D., Chen, Z. W. S., Jang, J., Shevchenko, A., Charbonneau, H., and Deshaies, R. J. (1999) Exit from Mitosis Is Triggered by Tem1-Dependent Release of the Protein Phosphatase Cdc14 from Nucleolar RENT Complex. Cell. 97, 233–244

13. Straight, A. F., Shou, W., Dowd, G. J., Turck, C. W., Deshaies, R. J., Johnson, A. D., and Moazed, D. (1999) Net1, a Sir2-Associated Nucleolar Protein Required for rDNA Silencing and Nucleolar Integrity. Cell. 97, 245–256

14. Li, C., Mueller, J. E., and Bryk, M. (2006) Sir2 represses endogenous polymerase II transcription units in the ribosomal DNA nontranscribed spacer. Molecular biology of the cell. 17, 3848–3859

15. Bryk, M., Banerjee, M., Murphy, M., Knudsen, K. E., Garfinkel, D. J., and Curcio, M. J. (1997) Transcriptional silencing of Ty1 elements in the RDN1 locus of yeast. Genes & Development. 11, 255–269

16. Smith, J. S., and Boeke, J. D. (1997) An unusual form of transcriptional silencing in yeast ribosomal DNA. Genes & Development. 11, 241–254

17. Gottlieb, S., and Esposito, R. E. (1989) A new role for a yeast transcriptional silencer gene, SIR2, in regulation of recombination in ribosomal DNA. Cell. 56, 771–776

18. Ganley, A. R. D., and Kobayashi, T. (2014) Ribosomal DNA and cellular senescence: new evidence supporting the connection between rDNA and aging. FEMS Yeast Research. 14, 49–59

19. Buck, S. W., Sandmeier, J. J., and Smith, J. S. (2002) RNA Polymerase I Propagates Unidirectional Spreading of rDNA Silent Chromatin. Cell. 111, 1003–1014

20. Kaeberlein, M., McVey, M., and Guarente, L. (1999) The SIR2/3/4 complex and SIR2 alone promote longevity in Saccharomyces cerevisiae by two different mechanisms. Genes Dev. 13, 2570–2580

21. Smith, J. S., Brachmann, C. B., Pillus, L., and Boeke, J. D. (1998) Distribution of a Limited Sir2 Protein Pool Regulates the Strength of Yeast rDNA Silencing and Is Modulated by Sir4p. Genetics. 149, 1205–1219

22. Sinclair, D. A., and Guarente, L. (1997) Extrachromosomal rDNA Circles— A Cause of Aging in Yeast. Cell. 91, 1033–1042

23. Johzuka, K., and Horiuchi, T. (2002) Replication fork block protein, Fob1, acts as an rDNA region specific recombinator in S. cerevisiae. Genes to Cells. 7, 99–113

24. Defossez, P.-A., Prusty, R., Kaeberlein, M., Lin, S.-J., Ferrigno, P., Silver, P. A., Keil, R. L., and Guarente, L. (1999) Elimination of Replication Block Protein Fob1 Extends the Life Span of Yeast Mother Cells. Molecular Cell. 3, 447–455

25. Hotz, M., Thayer, N. H., Hendrickson, D. G., Schinski, E. L., Xu, J., and Gottschling, D. E. (2022) rDNA array length is a major determinant of replicative lifespan in budding yeast. Proceedings of the National Academy of Sciences. 119, e2119593119

26. Hu, Z., Xia, B., Postnikoff, S. D., Shen, Z.-J., Tomoiaga, A. S., Harkness, T. A., Seol, J. H., Li, W., Chen, K., and Tyler, J. K. (2018) Ssd1 and Gcn2 suppress global translation efficiency in replicatively aged yeast while their activation extends lifespan. eLife. 7, e35551

27. Janssens, G. E., Meinema, A. C., González, J., Wolters, J. C., Schmidt, A., Guryev, V., Bischoff, R., Wit, E. C., Veenhoff, L. M., and Heinemann, M. (2015) Protein biogenesis machinery is a driver of replicative aging in yeast. eLife. 4, e08527

28. Paxman, J., Zhou, Z., O’Laughlin, R., Liu, Y., Li, Y., Tian, W., Su, H., Jiang, Y., Holness, S. E., Stasiowski, E., Tsimring, L. S., Pillus, L., Hasty, J., and Hao, N. (2022) Age-dependent aggregation of ribosomal RNA-binding proteins links deterioration in chromatin stability with challenges to proteostasis. eLife. 11, e75978

29. Dang, W., Steffen, K. K., Perry, R., Dorsey, J. A., Johnson, F. B., Shilatifard, A., Kaeberlein, M., Kennedy, B. K., and Berger, S. L. (2009) Histone H4 lysine 16 acetylation regulates cellular lifespan. Nature. 459, 802–807

30. Fine, R. D., Maqani, N., Li, M., Franck, E., and Smith, J. S. (2019) Depletion of Limiting rDNA Structural Complexes Triggers Chromosomal Instability and Replicative Aging of Saccharomyces cerevisiae. Genetics. 212, 75–91

31. Pal, S., Postnikoff, S. D., Chavez, M., and Tyler, J. K. (2018) Impaired cohesion and homologous recombination during replicative aging in budding yeast. Science Advances. 4, eaaq0236

32. Hendrickson, D. G., Soifer, I., Wranik, B. J., Kim, G., Robles, M., Gibney, P. A., and McIsaac, R. S. (2018) A new experimental platform facilitates assessment of the transcriptional and chromatin landscapes of aging yeast. eLife. 7, e39911

33. Erjavec, N., Larsson, L., Grantham, J., and Nyström, T. (2007) Accelerated aging and failure to segregate damaged proteins in Sir2 mutants can be suppressed by overproducing the protein aggregation-remodeling factor Hsp104p. Genes Dev. 21, 2410–2421

34. Kolberg, L., Raudvere, U., Kuzmin, I., Adler, P., Vilo, J., and Peterson, H. (2023) g:Profiler—interoperable web service for functional enrichment analysis and gene identifier mapping (2023 update). Nucleic Acids Research. 51, W207–W212

35. Hu, Z., Chen, K., Xia, Z., Chavez, M., Pal, S., Seol, J.-H., Chen, C.-C., Li, W., and Tyler, J. K. (2014) Nucleosome loss leads to global transcriptional up-regulation and genomic instability during yeast aging. Genes Dev. 28, 396–408

36. Sun, Y., Yu, R., Guo, H.-B., Qin, H., and Dang, W. (2021) A quantitative yeast aging proteomics analysis reveals novel aging regulators. GeroScience. 43, 2573–2593

37. Wang, J. C. (2002) Cellular roles of DNA topoisomerases: a molecular perspective. Nature Reviews Molecular Cell Biology. 3, 430–440

38. Di Felice, F., Egidi, A., D’Alfonso, A., and Camilloni, G. (2019) Fob1p recruits DNA topoisomerase I to ribosomal genes locus and contributes to its transcriptional silencing maintenance. The International Journal of Biochemistry & Cell Biology. 110, 143–148

39. D’Alfonso, A., Di Felice, F., Carlini, V., Wright, C. M., Hertz, M. I., Bjornsti, M.-A., and Camilloni, G. (2016) Molecular Mechanism of DNA Topoisomerase I-Dependent rDNA Silencing: Sir2p Recruitment at Ribosomal Genes. J Mol Biol. 428, 4905–4916

40. Gardner, R. G., Nelson, Z. W., and Gottschling, D. E. (2005) Degradation-Mediated Protein Quality Control in the Nucleus. Cell. 120, 803–815

41. Arita, Y., Kim, G., Li, Z., Friesen, H., Turco, G., Wang, R. Y., Climie, D., Usaj, M., Hotz, M., Stoops, E. H., Baryshnikova, A., Boone, C., Botstein, D., Andrews, B. J., and McIsaac, R. S. (2021) A genome-scale yeast library with inducible expression of individual genes. Molecular Systems Biology. 17, e10207

42. Sloan, R., Huang, S. N., Pommier, Y., and Jinks-Robertson, S. (2017) Effects of camptothecin or TOP1 overexpression on genetic stability in Saccharomyces cerevisiae. DNA Repair. 59, 69–75

43. Buck, S. W., Maqani, N., Matecic, M., Hontz, R. D., Fine, R. D., Li, M., and Smith, J. S. (2016) RNA Polymerase I and Fob1 contributions to transcriptional silencing at the yeast rDNA locus. Nucleic Acids Research. 44, 6173–6184

44. Moreno, D. F., Jenkins, K., Morlot, S., Charvin, G., Csikasz-Nagy, A., and Aldea, M. (2019) Proteostasis collapse, a hallmark of aging, hinders the chaperone-Start network and arrests cells in G1. eLife. 8, e48240

45. Lin, S. S., Manchester, J. K., and Gordon, J. I. (2001) Enhanced Gluconeogenesis and Increased Energy Storage as Hallmarks of Aging in Saccharomyces cerevisiae * 210. Journal of Biological Chemistry. 276, 36000–36007

46. van der Knaap, J. A., and Verrijzer, C. P. (2025) Moonlighting Enzymes at the Interface Between Metabolism and Epigenetics. Annu Rev Biochem. 94, 279–303

47. Choudhary, R., Niska-Blakie, J., Adhil, M., Liberi, G., Achar, Y. J., Giannattasio, M., and Foiani, M. (2023) Sen1 and Rrm3 ensure permissive topological conditions for replication termination. Cell Reports. 42, 112747

48. Ivessa, A. S., Zhou, J.-Q., and Zakian, V. A. (2000) The Saccharomyces Pif1p DNA Helicase and the Highly Related Rrm3p Have Opposite Effects on Replication Fork Progression in Ribosomal DNA. Cell. 100, 479–489

49. Lanz, M. C., Zhang, S., Swaffer, M. P., Ziv, I., Götz, L. H., Kim, J., McCarthy, F., Jarosz, D. F., Elias, J. E., and Skotheim, J. M. (2024) Genome dilution by cell growth drives starvation-like proteome remodeling in mammalian and yeast cells. Nat Struct Mol Biol. 31, 1859–1871

50. Christiano, R., Nagaraj, N., Fröhlich, F., and Walther, T. C. (2014) Global Proteome Turnover Analyses of the Yeasts S. cerevisiae and S. pombe. Cell Reports. 9, 1959–1965

51. Zhao, T., Chida, A., Shichino, Y., Choi, D., Mizunuma, M., Iwasaki, S., and Ohya, Y. (2022) Multifarious Translational Regulation during Replicative Aging in Yeast. Journal of Fungi. 10.3390/jof8090938

52. Jazwinski, S. M., Egilmez, N. K., and Chen, J. B. (1989) Replication control and cellular life span. Exp Gerontol. 24, 423–436

53. Dirick, L., Böhm, T., and Nasmyth, K. (1995) Roles and regulation of Cln-Cdc28 kinases at the start of the cell cycle of Saccharomyces cerevisiae. EMBO J. 14, 4803–4813

54. Pérez-Ortín, J. E., Mena, A., Barba-Aliaga, M., Singh, A., Chávez, S., and García-Martínez, J. (2021) Cell volume homeostatically controls the rDNA repeat copy number and rRNA synthesis rate in yeast. PLoS Genet. 17, e1009520

55. Christman, M. F., Dietrich, F. S., and Fink, G. R. (1988) Mitotic recombination in the rDNA of S. cerevisiae is suppressed by the combined action of DNA topoisomerases I and II. Cell. 55, 413–425

56. Kim, R. A., and Wang, J. C. (1989) A subthreshold level of DNA topoisomerases leads to the excision of yeast rDNA as extrachromosomal rings. Cell. 57, 975–985

57. Cioci, F., Vogelauer, M., and Camilloni, G. (2002) Acetylation and Accessibility of rDNA Chromatin in Saccharomyces cerevisiae in Δtop1 and Δsir2 Mutants. Journal of Molecular Biology. 322, 41–52

58. Andersen, S. L., Sloan, R. S., Petes, T. D., and Jinks-Robertson, S. (2015) Genome-Destabilizing Effects Associated with Top1 Loss or Accumulation of Top1 Cleavage Complexes in Yeast. PLOS Genetics. 11, e1005098

59. El Hage, A., French, S. L., Beyer, A. L., and Tollervey, D. (2010) Loss of Topoisomerase I leads to R-loop-mediated transcriptional blocks during ribosomal RNA synthesis. Genes & Development. 24, 1546–1558

60. Smith, J. S., Caputo, E., and Boeke, J. D. (1999) A genetic screen for ribosomal DNA silencing defects identifies multiple DNA replication and chromatin-modulating factors. Mol Cell Biol. 19, 3184–3197

61. Schultz, M. C., Brill, S. J., Ju, Q., Sternglanz, R., and Reeder, R. H. (1992) Topoisomerases and yeast rRNA transcription: negative supercoiling stimulates initiation and topoisomerase activity is required for elongation. Genes Dev. 6, 1332–1341

62. Liu, L. F., and Wang, J. C. (1987) Supercoiling of the DNA template during transcription. Proc. Natl. Acad. Sci. U.S.A. 84, 7024–7027

63. Cheng, T. H., Li, Y. C., and Gartenberg, M. R. (1998) Persistence of an alternate chromatin structure at silenced loci in the absence of silencers. Proc Natl Acad Sci U S A. 95, 5521– 5526

64. Matecic, M., Smith, D. L., Pan, X., Maqani, N., Bekiranov, S., Boeke, J. D., and Smith, J. S. (2010) A microarray-based genetic screen for yeast chronological aging factors. PLoS Genet. 6, e1000921

65. Andrews, B., Boone, C. M., Davis, T., Fields, S., and Cold Spring Harbor Laboratory (eds.) (2016) Budding yeast: a laboratory manual, Cold Spring Harbor Laboratory Press, Cold Spring Harbor, New York

66. Longtine, M. S., Mckenzie III, A., Demarini, D. J., Shah, N. G., Wach, A., Brachat, A., Philippsen, P., and Pringle, J. R. (1998) Additional modules for versatile and economical PCR-based gene deletion and modification in Saccharomyces cerevisiae. Yeast. 14, 953– 961

67. Colley, W. C., van der Merwe, M., Vance, J. R., Burgin, A. B., Jr., and Bjornsti, M.-A. (2004) Substitution of Conserved Residues within the Active Site Alters the Cleavage Religation Equilibrium of DNA Topoisomerase I *. Journal of Biological Chemistry. 279, 54069–54078

68. Isasa, M., Rose, C. M., Elsasser, S., Navarrete-Perea, J., Paulo, J. A., Finley, D. J., and Gygi, S. P. (2015) Multiplexed, Proteome-Wide Protein Expression Profiling: Yeast Deubiquitylating Enzyme Knockout Strains. J. Proteome Res. 14, 5306–5317

69. Miller, A. W., Befort, C., Kerr, E. O., and Dunham, M. J. (2013) Design and Use of Multiplexed Chemostat Arrays. JoVE. 10.3791/50262-v

70. Collart, M. A., and Oliviero, S. (1993) Preparation of Yeast RNA. Current Protocols in Molecular Biology. 23, 13.12.1–13.12.5

71. Dunn, B., and Wobbe, C. R. (1993) Preparation of Protein Extracts from Yeast. Current Protocols in Molecular Biology. 23, 13.13.1–13.13.9

72. Buchanan, B. W., Lloyd, M. E., Engle, S. M., and Rubenstein, E. M. (2016) Cycloheximide Chase Analysis of Protein Degradation in Saccharomyces cerevisiae. JoVE. 10.3791/53975

73. Steffen, K. K., Kennedy, B. K., and Kaeberlein, M. (2009) Measuring Replicative Life Span in the Budding Yeast. JoVE. 10.3791/1209

74. Jo, M. C., Liu, W., Gu, L., Dang, W., and Qin, L. (2015) High-throughput analysis of yeast replicative aging using a microfluidic system. Proceedings of the National Academy of Sciences. 112, 9364–9369

75. Han, S. K., Lee, D., Lee, H., Kim, D., Son, H. G., Yang, J.-S., Lee, S.-J. V., and Kim, S. (2016) OASIS 2: online application for survival analysis 2 with features for the analysis of maximal lifespan and healthspan in aging research. Oncotarget; Vol 7, No 35. [online] https://www.oncotarget.com/article/11269/ (Accessed January 1, 2016)

76. Li, M., Petteys, B. J., McClure, J. M., Valsakumar, V., Bekiranov, S., Frank, E. L., and Smith, J. S. (2010) Thiamine biosynthesis in Saccharomyces cerevisiae is regulated by the NAD+-dependent histone deacetylase Hst1. Mol Cell Biol. 30, 3329–3341

77. Borten, M. A., Bajikar, S. S., Sasaki, N., Clevers, H., and Janes, K. A. (2018) Automated brightfield morphometry of 3D organoid populations by OrganoSeg. Sci Rep. 8, 5319

78. Lindstrom, D. L., Leverich, C. K., Henderson, K. A., and Gottschling, D. E. (2011) Replicative age induces mitotic recombination in the ribosomal RNA gene cluster of Saccharomyces cerevisiae. PLoS Genet. 7, e1002015

79. Baker Brachmann, C., Davies, A., Cost, G. J., Caputo, E., Li, J., Hieter, P., and Boeke, J. D. (1998) Designer deletion strains derived from Saccharomyces cerevisiae S288C: A useful set of strains and plasmids for PCR-mediated gene disruption and other applications. Yeast. 14, 115–132

80. Sikorski, R. S., and Hieter, P. (1989) A system of shuttle vectors and yeast host strains designed for efficient manipulation of DNA in Saccharomyces cerevisiae. Genetics. 122, 19–27

81. Chu, A. M., and Davis, R. W. (2008) High-Throughput Creation of a Whole-Genome Collection of Yeast Knockout Strains. in Microbial Gene Essentiality: Protocols and Bioinformatics (Osterman, A. L., and Gerdes, S. Y. eds), pp. 205–220, Methods in Molecular BiologyTM, Humana Press, Totowa, NJ, 416, 205–220

