## Supporting Information for "Age-dependent topoisomerase I depletion alters recruitment of rDNA silencing complexes"

**Figures S1-S5**

**Tables S1-S4**

**Excel File:**

**Table S5.** Complete proteomic dataset comparing protein levels of isolated nuclei from replicatively aged (~6-7 generations) and young (~0-2 generations) yeast cells using TMT-MS.

**Figure S1**. **(A)** Representative image of DAPI stained nuclei isolated from strain SY38. **(B)** Histone acetylase (HAT) activity assay performed on isolated nuclei from young and old cells, and a positive control using HeLa cell nuclear extract (4 mg/ml). ****$p<0.0001$ (one-way ANOVA with Dunnett's correction for multiple comparisons compared to positive control, n=5).

A

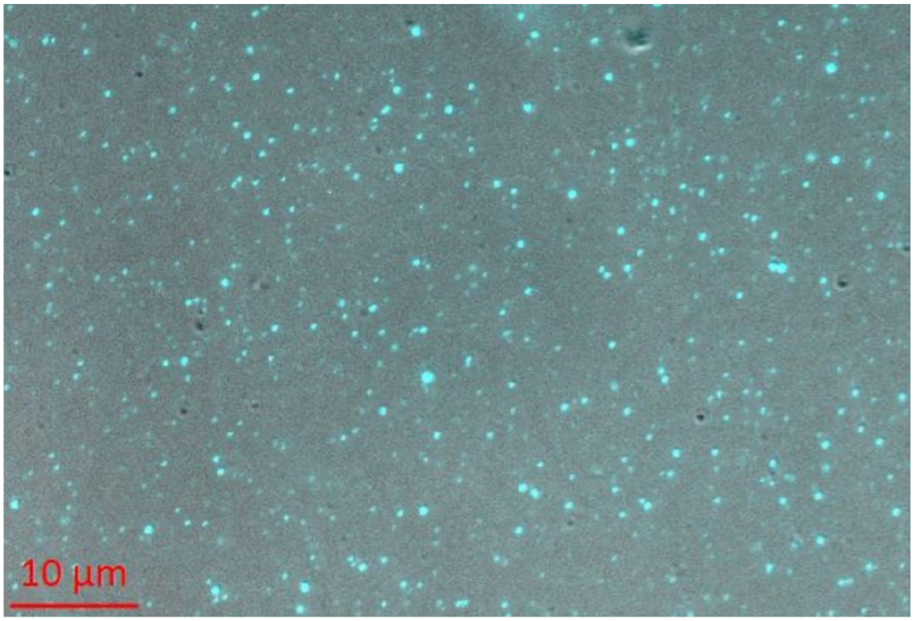
10 µm

B

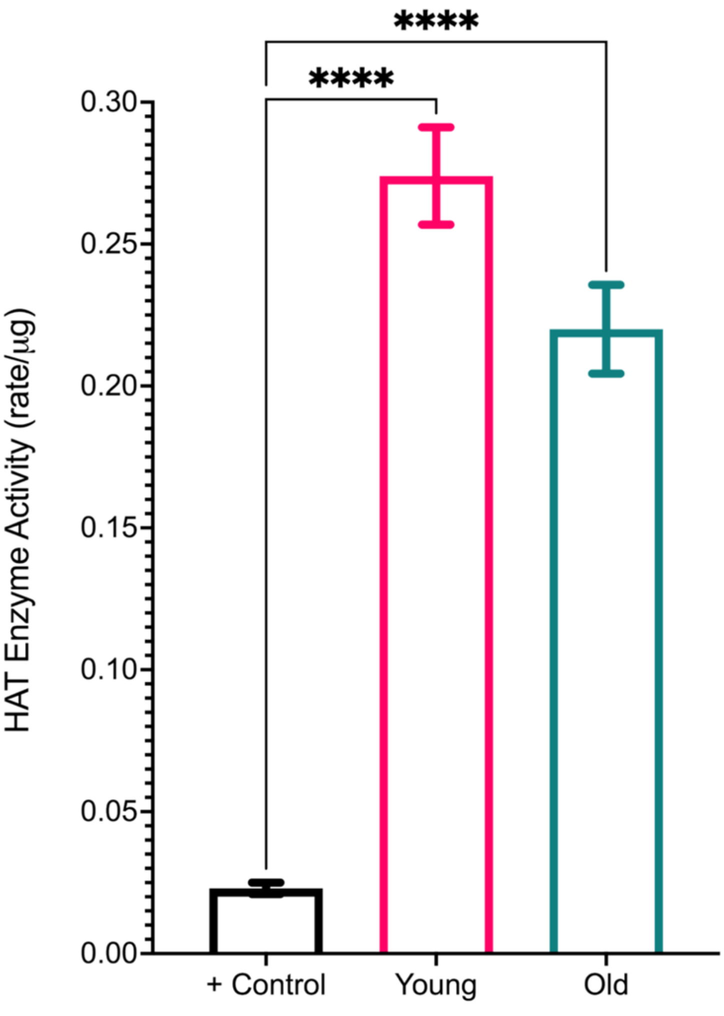
****
****
0.30
0.25
0.20
0.15
0.10
0.05
0.00
HAT Enzyme Activity (rate/µg)
+ Control
Young
Old

**Figure S2**. Quantitative RT-PCR of lncRNA expression from the rDNA IGS1. The fold change relative to WT (BY4741) is indicated for Top1-Myc tagged (LP128) or *sir2*Δ (SY533) strains. (One-way ANOVA with Dunnett's test for multiple comparisons, p= 0.9812, *p=0.0171, n=3).

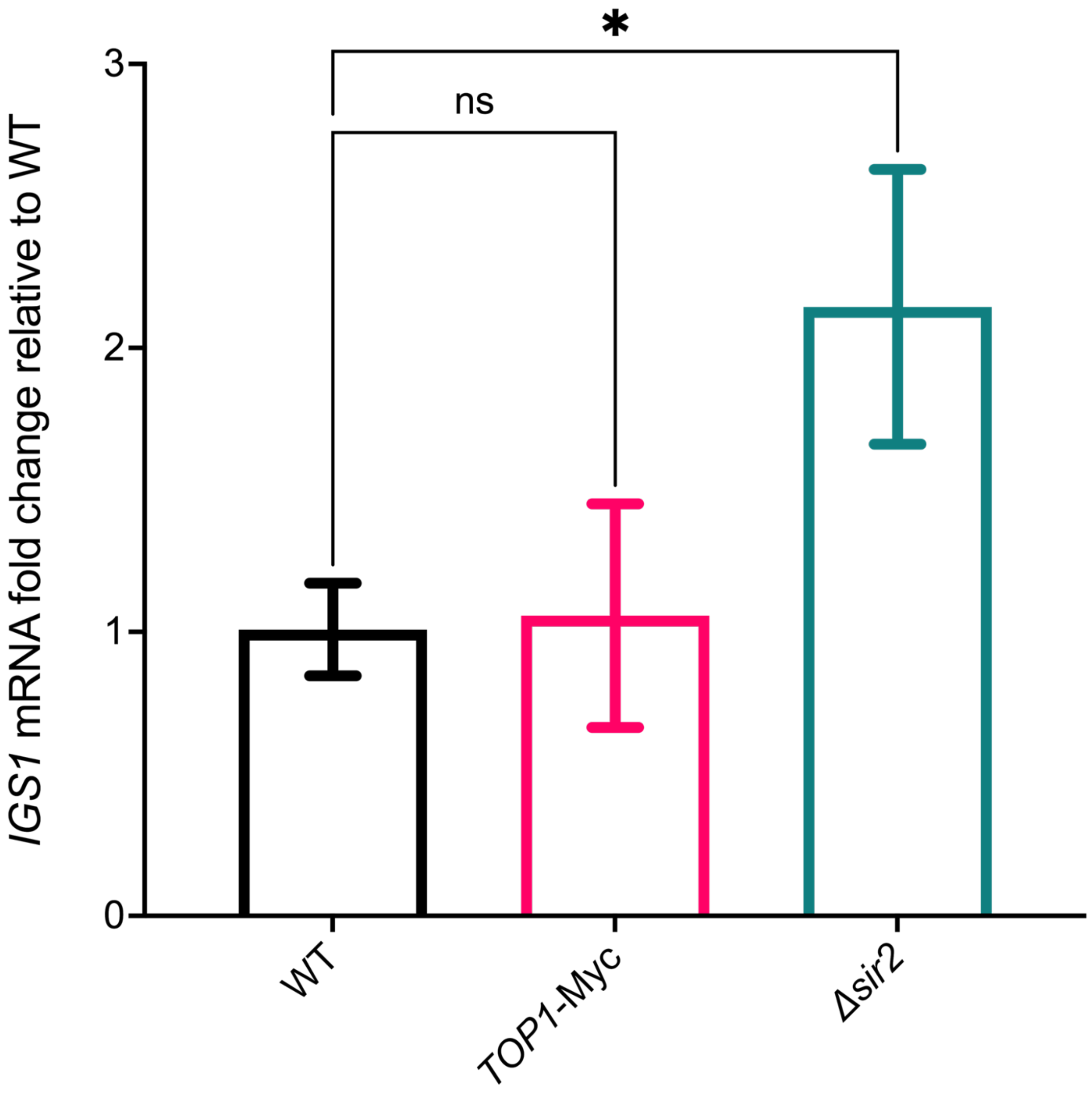

**Figure S3**. Cycloheximide chase experiment validation. **(A)** Growth curve measured by $OD_{600}$ of wildtype yeast strain BY4741 grown in liquid YPD media with a final concentration of 0, 125, 250, 250, or 500 μg/mL cycloheximide. **(B)** Cdc13-13xMyc protein levels after 30, 60, 90, 120, or 240 minutes of growth after addition of 250 μg/mL cycloheximide at non-permissive temperature (30°C). GAPDH is used as the loading control.

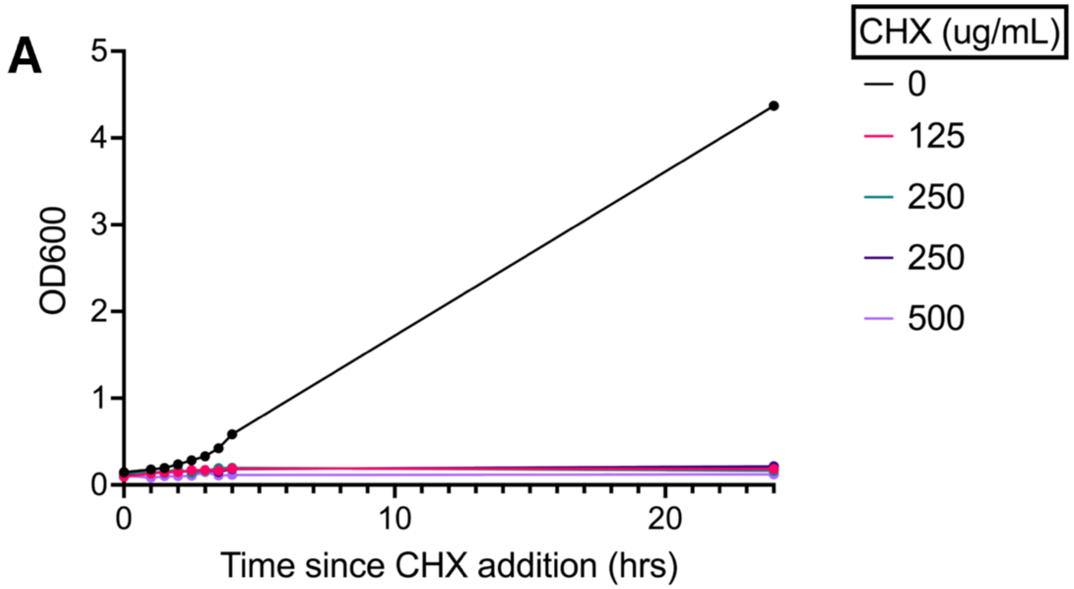
A
5
4
3
2
1
0
OD600
0
10
20
Time since CHX addition (hrs)
CHX (ug/mL)
0
125
250
250
500

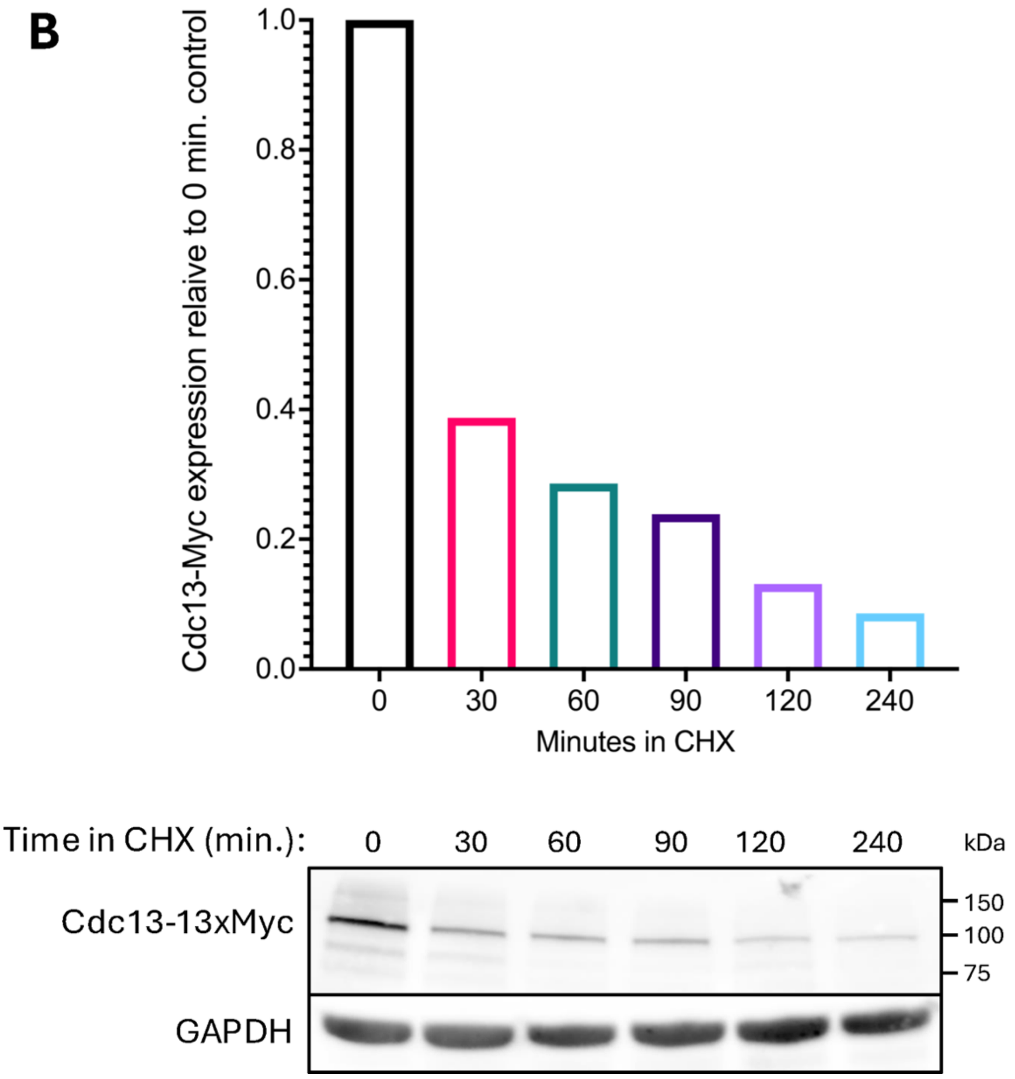
B
1.0
0.8
0.6
0.4
0.2
0.0
Cdc13-Myc expression relaive to 0 min. control
0
30
60
90
120
240
Minutes in CHX
Time in CHX (min.):
0
30
60
90
120
240
kDa
Cdc13-13xMyc
150
100
75
GAPDH

**Figure S4**. Top1-Myc expression from the YETI overexpression strain LP124 induced with 0, 2.5, or 5 nM estradiol as compared to the WT control (Border Strain LP55) (one-way ANOVA with Dunnett's test for multiple comparisons, **p=0.0027, ***p=0.0005, **p=0.0035 n=3). Tub1 is used as the loading control.

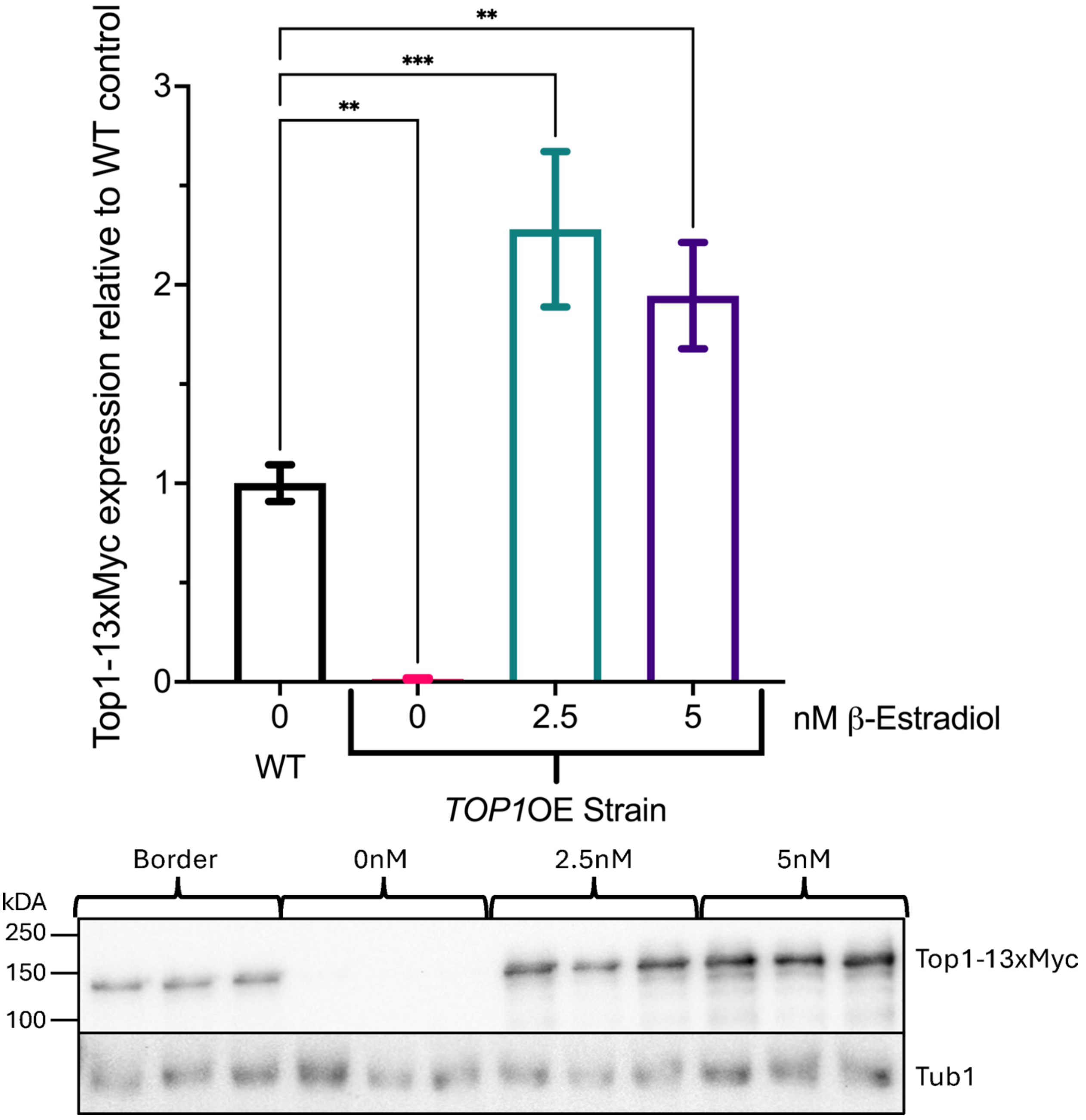

**Figure S5**. Verification by qRT-PCR of *TOP1* mRNA overexpression by low copy (CEN/ARS) or high copy (2µ) *LEU2* vectors expressing either wildtype (*TOP1*-WT) or catalytically dead (*top1*-CD) under control of the native *TOP1* promoter. Fold change is relative to a strain harboring the empty vector control. (CEN/ARS: **p=0.0014, **p=0.0037, one-way ANOVA with Dunnett's test for multiple-comparisons, n=3) (2µ: **p=0.0023, *p=0.0325 one-way ANOVA with Dunnett's test for multiple-comparisons, n=3).

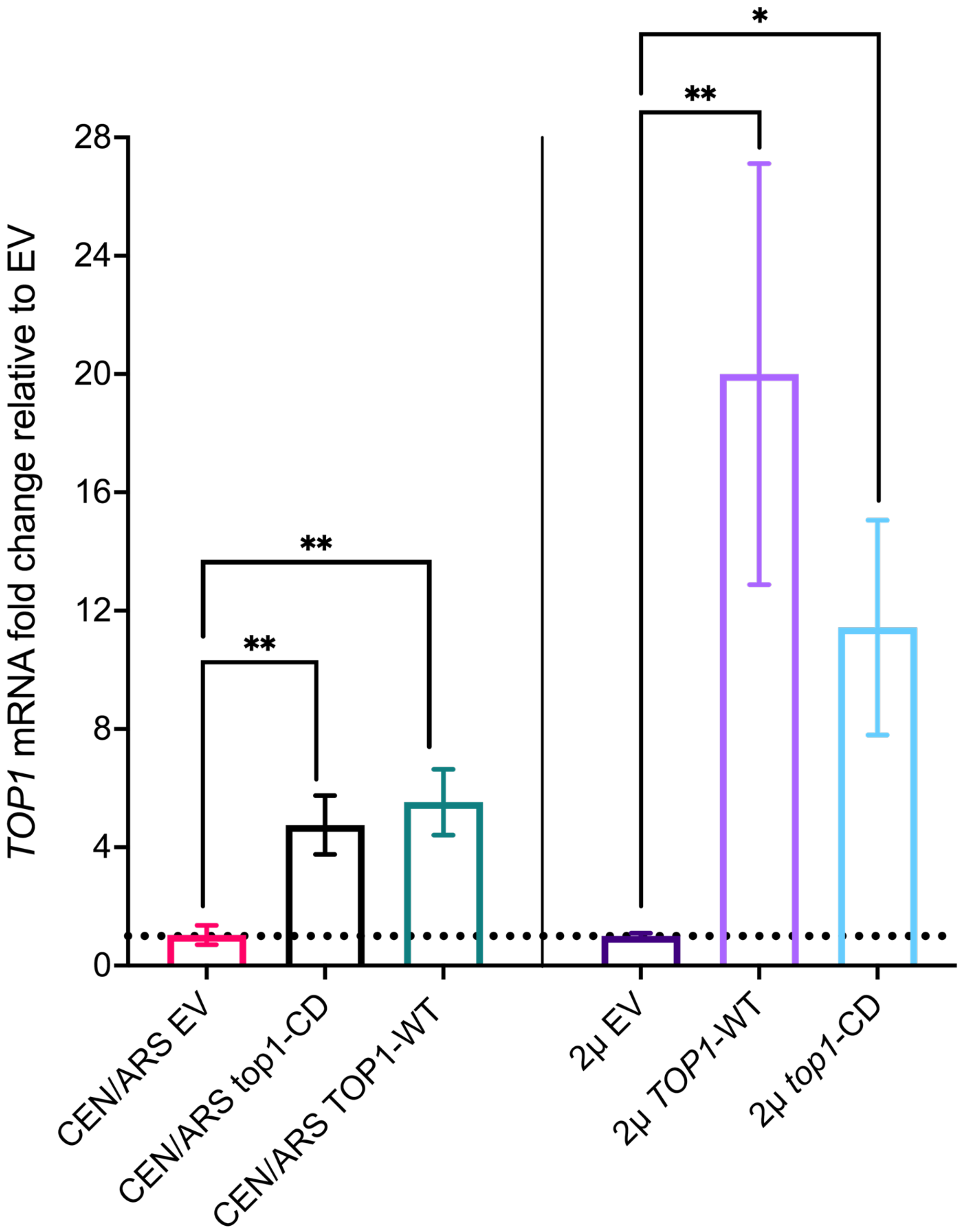

TOP1 mRNA fold change relative to EV
0
4
8
12
16
20
24
28
**
**
**
*
CEN/ARS EV
CEN/ARS top1-CD
CEN/ARS TOP1-WT
2μ EV
2μ TOP1-WT
2μ top1-CD

**Table S1. List of yeast strains.**

| Strain | Genotype | Source |
|---|---|---|
| BY4743 (SY38) | *MAT a his3$\Delta$1 leu2$\Delta$0 LYS2 met15$\Delta$0 ura3$\Delta$0/ MAT$\alpha$ his3$\Delta$1 leu2$\Delta$0 lys2$\Delta$0 MET15 ura3$\Delta$0* | (79) |
| BY4741 (SY41) | *MAT a his3$\Delta$1 leu2$\Delta$0 LYS2 met15$\Delta$0 ura3$\Delta$0* | (79) |
| YPH499 (SY104) | *MAT a ura3-52 lys2-801 ade2-101 trp1-$\Delta$63 his3-$\Delta$200 leu2-$\Delta$1* | (80) |
| MD188 | *MAT a his3$\Delta$1 leu2$\Delta$0 LYS2 met15$\Delta$0 ura3$\Delta$0 Hsp104-13xMyc::KanMX* | This study |
| YRH919 | *MAT $\alpha$ his3$\Delta$200 leu2$\Delta$1 trp1$\Delta$63 ura3-167 nts1$\Delta$::URA3/HIS3 Top1-13xMyc::KanMX* | This study |
| LP128 | *MAT a his3$\Delta$1 leu2$\Delta$0 LYS2 met15$\Delta$0 ura3$\Delta$0 Top1-13xMyc::KanMX* | This Study |
| LP55 | *MAT a [HAP1::natMX::ACT1pr-Z3EV-ENO2term] ura3$\Delta$0 can1$\Delta$::STE2pr-Sphis5 his3$\Delta$1 lyp1$\Delta$* | (41) |
| LP56 | *MAT a [barcode::URA3::Z3EVpr-Top1] [HAP1::natMX::ACT1pr-Z3EV-ENO2term] ura3$\Delta$0 can1$\Delta$::STE2pr-Sphis5 his3$\Delta$1 lyp1$\Delta$* | (41) |
| LP181 | *MAT a ura352 lys2-801 ade2-101 trp1$\Delta$63 his3$\Delta$200 leu2$\Delta$1 + pRS416* | This study |
| LP183 | *MAT a ura352 lys2-801 ade2-101 trp1$\Delta$63 his3$\Delta$200 leu2$\Delta$1 + pNK66 (YCpSCTOP1-U)* | This study |
| LP185 | *MAT a ura352 lys2-801 ade2-101 trp1$\Delta$63 his3$\Delta$200 leu2$\Delta$1 + pNK67 (YCpSCTOP1 Y727F-U)* | This study |
| LP257 | *MAT a ura352 lys2-801 ade2-101 trp1$\Delta$63 his3$\Delta$200 leu2$\Delta$1::SIR2-LEU2 + pRS416* | This study |
| LP263 | *MAT a ura352 lys2-801 ade2-101 trp1$\Delta$63 his3$\Delta$200 leu2$\Delta$1::SIR2-LEU2 + pNK66 (YCpSCTOP1-U)* | This study |
| LP269 | *MAT a ura352 lys2-801 ade2-101 trp1$\Delta$63 his3$\Delta$200 leu2$\Delta$1::SIR2-LEU2 + pNK67 (YCpSCTOP1 Y727F-U)* | This study |
| YNM44 | *MAT $\alpha$ his3$\Delta$200 leu2$\Delta$1 ura3-167 RDN1 (Ter1-R)::mURA3-HIS3* | (43) |
| LP301 | YNM44 *$\Delta$top1::KanMX* | This study |
| AW1 | YNM44 + pRS415 | This study |
| AW2 | YNM44 + pAW2 | This study |
| AW3 | YNM44 + pAW4 | This study |
| AW4 | YNM44 + pSB794 | This study |
| AW5 | YNM44 + pSB764 | This study |
| AW6 | YNM44 + pRS425 | This study |
| AW7 | YNM44 + pAW6 | This study |
| AW8 | YNM44 + pAW8 | This study |
| AW9 | YNM44 + pSB790 | This study |

| | | |
|---|---|---|
| AW10 | YNM44 + pSB766 | This study |
| AW11 | LP301 + pRS415 | This study |
| AW12 | LP301 + pAW2 | This study |
| AW13 | LP301 + pAW4 | This study |
| AW14 | LP301 + pSB794 | This study |
| AW15 | LP301 + pSB764 | This study |
| AW16 | LP301 + pRS425 | This study |
| AW17 | LP301 + pAW6 | This study |
| AW18 | LP301 + pAW8 | This study |
| AW19 | LP301 + pSB790 | This study |
| AW20 | LP301 + pSB766 | This study |
| AW41 | JS124 + pRS415 | This study |
| AW42 | JS124 + pAW2 | This study |
| AW43 | JS124 + pAW4 | This study |
| AW44 | JS124 + pSB794 | This study |
| AW45 | JS124 + pSB764 | This study |
| AW46 | JS124 + pRS425 | This study |
| AW47 | JS124 + pAW6 | This study |
| AW48 | JS124 + pAW8 | This study |
| AW49 | JS124 + pSB790 | This study |
| AW50 | JS124 + pSB766 | This study |
| MD207 | YPH499 *SIR2-13xMyc::KanMX* | This study |
| MD208 | MD207 + pRS416 | This study |
| MD209 | MD207 + pNK66 (*YCpSCTOP1-U*) | This study |
| MD210 | MD207 + pNK67 (*YCpSCtop1Y727F-U*) | This study |
| SY533 | BY4741 Δsir2::URA3 | (81) |
| LP124 | LP56 *TOP1-13xMyc::KanMX* | This Study |
| UCC6277 (RGY525) | *MATa ade2Δ::hisG his3Δ200 leu2Δ0 lys2Δ0 met15Δ0 trp1Δ63 ura3Δ0 ADE2-TEL-VR sir4::hphMX::pTDH3-1Myc-sir4-9::LEU2 CDC13::TRP1::cdc13-1-9Myc* | (40) |

**Table S2**. **List of plasmids**.

| Plasmid | Description | Source |
|---|---|---|
| pNK66 (YCpSc*TOP1*-U) | WT *TOP1* under its native promoter in a pRS416 yeast CEN/ARS vector. | (67) (Gifted by Nyun Kim.) |
| pNK67 (YCpSc*top1* Y727F-U) | Catalytically dead Top1 under its native promoter in pRS416 yeast CEN/ARS vector. | (67) (Gifted by Nyun Kim.) |
| pAsc415 | CEN/ARS *LEU2* (*Asc*I site ligated into *SmaI* site of pRS415) | (19) |
| pSB794 | pAsc415-*SIR2* | (19) |
| pSB764 | pAsc415-*NET1* | (19) |
| pAW4 | pAsc415-*TOP1* (WT) | This Paper |
| pAW2 | pAsc415-*top1*Y727F (CD) | This Paper |
| pAsc425 | *2μ LEU2* (*Asc*I site ligated into *SmaI* site of pRS425) | (19) |
| pSB766 | pAsc425-*SIR2* | (19) |
| pSB790 | pAsc425-*NET1* | (19) |
| pAW6 | pAsc425-*TOP1* (WT) | This Paper |
| pAW8 | pAsc425-*top1*Y727F (CD) | This Paper |

**Table S3**. **List of DNA oligos**.

| Oligo | Description | Sequence 5'-3' |
|---|---|---|
| JS3878 | *HSP104-13xMyc* Fw | CGATAATGAGGACAGTATGGAAATTGATGATGACC TAGATCGGATCCCCGGGTTAATTAA |
| JS3879 | *HSP104-13xMyc* Rv | ATTCTTGTTCGAAAGTTTTTAAAAATCACACTATAT TAAAGAATTCGAGCTCGTTTAAAC |
| JS3611 | *TOP1-13xMyc* Fw | GTTCCGATTGAAAAGATTTT |
| JS3612 | *TOP1-13xMyc* Rv | CTTCCTAGTAACCCTAATGC |
| JS3398 | RTqPCR *TOP1* Fw | CGAGAAGAAGAAGAAGAGGAGG |
| JS3399 | RTqPCR *TOP1* Rv | TGGTAAGGGCTGGTATGGTG |
| JS3976 | RTqPCR *UBC6* Fw | ATTGGATGAGGGGGATGCGGCA |
| JS3977 | RTqPCR *UBC6* Rv | AGCGCGTATTCTGTCTTCAGGGT |
| JS4054 | *TOP1* short gene PCR cloning (AscI) Fw | TAGGCGCGCCATATGATCGATGCACGTAAAGAAC |
| JS4055 | *TOP1* short gene PCR cloning (AscI) Rv | TAGGCGCGCCAAGAGATACGGACAATACGTTTC |
| JS1191 | *SIR2-13xMyc* Fw | CGTGTATGTCGTTACATCAGATGAACATCCCAAAA CCCTCCGGATCCCCGGGTTAATTAA |
| JS1192 | *SIR2-13xMyc* Rv | TATTAATTTGGCACTTTTAAATTATTAAATTGCCTT CTACGAATTCGAGCTCGTTTAAAC |

**Table S4. CHX Chase Assay P-Value reporting.**

| Dunnett's multiple comparisons test | Mean Diff. | 95.00% CI of diff. | Below threshold? | Summary | Adjusted P Value |
|---|---|---|---|---|---|
| **Figure 2C** | | | | | |
| 0 vs. 30 | 0.2355 | -0.3085 to 0.7795 | No | ns | 0.4948 |
| 0 vs. 60 | 0.09269 | -0.4513 to 0.6367 | No | ns | 0.9264 |
| 0 vs. 90 | 0.3247 | -0.2193 to 0.8687 | No | ns | 0.2712 |
| **Figure 2D** | | | | | |
| 0 vs. 1 | -0.1274 | -1.165 to 0.9098 | No | ns | 0.9694 |
| 0 vs. 2 | -0.219 | -1.256 to 0.8182 | No | ns | 0.8746 |
| 0 vs. 4 | -0.01247 | -1.050 to 1.025 | No | ns | >0.9999 |
| **Figure 2E (0-2 buds)** | | | | | |
| 0 vs. 30 | -0.0736 | -0.6281 to 0.4809 | No | ns | 0.9622 |
| 0 vs. 60 | -0.0304 | -0.5849 to 0.5241 | No | ns | 0.997 |
| 0 vs. 90 | 0.09023 | -0.4643 to 0.6448 | No | ns | 0.9347 |
| **Figure 2E (6-7 buds)** | | | | | |
| 0 vs. 30 | -0.2456 | -1.131 to 0.6397 | No | ns | 0.7688 |
| 0 vs. 60 | -0.0722 | -0.9575 to 0.8131 | No | ns | 0.9904 |
| 0 vs. 90 | -0.03633 | -0.9216 to 0.8489 | No | ns | 0.9987 |
